# Biallelic variants in the RNA exosome gene *EXOSC5* are associated with developmental delays, short stature, cerebellar hypoplasia and motor weakness

**DOI:** 10.1101/2020.04.01.839274

**Authors:** Anne Slavotinek, Doriana Misceo, Stephanie Htun, Linda Mathisen, Eirik Frengen, Michelle Foreman, Jennifer E. Hurtig, Liz Enyenihi, Maria C. Sterrett, Sara W. Leung, Dina Schneidman-Duhovny, Juvianee Estrada-Veras, Jacque L. Duncan, Vivian Xia, Daniah Beleford, Yue Si, Ganka Douglas, Hans Einar Treidene, Ambro van Hoof, Milo B. Fasken, Anita H. Corbett

**Author notes:** These two authors contributed equally to this work (Anne Slavotinek, Doriana Misceo). These two authors contributed equally to this work (Milo B. Fasken, Anita H. Corbett).

## Abstract

The RNA exosome is an essential ribonuclease complex involved in the processing and degradation of both coding and noncoding RNAs. We present three patients with biallelic variants in *EXOSC5*, which encodes a structural subunit of the RNA exosome. The common clinical features of these patients comprise failure to thrive, short stature, feeding difficulties, developmental delays that affect motor skills, hypotonia and esotropia. Brain MRI revealed cerebellar hypoplasia and ventriculomegaly. The first patient had a deletion involving exons 5-6 of *EXOSC5* and a missense variant, p.Thr114Ile, that were inherited *in trans*, the second patient was homozygous for p.Leu206His, and the third patient had paternal isodisomy for chromosome 19 and was homozygous for p.Met148Thr. We employed three complementary approaches to explore the requirement for *EXOSC5* in brain development and assess the functional consequences of pathogenic variants in *EXOSC5*. Loss of function for the zebrafish ortholog results in shortened and curved tails and bodies, reduced eye and head size and edema. We modeled pathogenic *EXOSC5* variants in both budding yeast and mammalian cells. Some of these variants show defects in RNA exosome function as well as altered interactions with other RNA exosome subunits. Overall, these findings expand the number of genes encoding RNA exosome components that have been implicated in human disease, while also suggesting that disease mechanism varies depending on the specific pathogenic variant.

## Introduction

The RNA exosome is a ribonuclease complex composed of ten, evolutionarily conserved subunits that form a ring-like structure with 3’-5’ exonuclease activity that is critical for the processing and degradation of a variety of RNAs both in the nucleus and cytoplasm (1-3). The exosome subunits are designated as exosome component (EXOSC) proteins in humans, mice (*M. musculus*) and other mammals and ribosomal RNA processing (Rrp) proteins in zebrafish (*D. rerio*), fruit flies (*D. melanogaster*) and budding yeast (*S. cerevisiae*), where the complex was first identified and studied (4, 5). The EXOSC4-9 subunits and their orthologs form a central, six-subunit ring that is covered by a three-subunit cap composed of EXOSC1-3. The ring-like complex creates a central channel through which RNA substrates are threaded to the catalytic subunit, DIS3 or DIS3L, at the base (6-10). In addition to DIS3 or DIS3L, the 9-subunit exosome complex also associates with another catalytic subunit, EXOSC10 at the cap. Notably, DIS3 and EXOSC10 are predominantly nuclear (7, 9-11), whereas DIS3L is mostly cytoplasmic (12). Although all six structural core ring subunits contain an RNase PH-like domain, they are all catalytically inactive due to amino acid substitutions that replace key catalytic residues (13). Thus, RNA substrates of the RNA exosome access the DIS3 catalytic subunit primarily via interactions with the non-catalytic core channel (14-16). Consistent with the critical function of the RNA exosome in the processing and decay of numerous cellular RNAs, all structural subunits and the DIS3 catalytic base are essential in all organisms analyzed thus far (3, 17).

The critical and conserved role of the RNA exosome implies that variants that alter the function of this complex could result in human disease. Indeed, deleterious variants in four genes – *EXOSC2, EXOSC3, EXOSC8*, and *EXOSC9* - encoding structural subunits of the RNA exosome and in several exosome cofactors have so far been linked to clinical phenotypes (Table 1) (18). Biallelic variants in *EXOSC3*, encoding a cap exosome subunit, cause pontocerebellar hypoplasia type 1b (PCH1b, OMIM 614678), a neurodegenerative condition characterized by neonatal hypotonia, feeding and swallowing difficulties, respiratory insufficiency, variable developmental delays, ataxia, lower motor neuron signs with muscle weakness and atrophy and limited survival (3, 19-28). Less common features, including seizures, nystagmus, strabismus and optic atrophy and dyskinesia, dystonia and spastic paraplegia, can be observed in patients with longer survival (24). Brain magnetic resonance imaging (MRI) findings revealed hypoplasia and/or atrophy of the cerebellum and pons, intra-cerebellar cysts and microcephaly (24). Biallelic variants in *EXOSC8*, encoding a ring exosome subunit, cause pontocerebellar hypoplasia type 1c (PCH1c, OMIM 616081), a very rare (only three pedigrees reported to date) disease with phenotypes that overlap with those in patients with *EXOSC3* variants and characterized by failure to thrive, respiratory distress, severe muscle weakness and signs of spinal muscular atrophy (SMA), spasticity, developmental delays and impairment of vision and hearing with onset between two to four months of age (29). Deterioration in clinical status can be triggered by intercurrent infections (29). Brain MRIs demonstrated cerebellar vermis hypoplasia, immature myelination and hypomyelination, cortical atrophy and thinning of the corpus callosum (29).

**Table 1.** Comparison of clinical features associated with mutations in RNA exosome subunit genes.

| **Clinical features** | **EXOSC3** | **EXOSC8** | **EXOSC2** | **EXOSC9** | **EXOSC5** |
| --- | --- | --- | --- | --- | --- |
| <u>Growth parameters</u> |  |  |  |  |  |
| Short stature/failure to thrive |  | + | + |  | 3/3 |
| Microcephaly | + |  |  |  | - |
| <u>Development</u> |  |  |  |  |  |
| Developmental delays | + | + |  | + | 3/3 |
| <u>Craniofacial anomalies</u> |  |  |  |  |  |
| Sparse and fine hair |  |  | + |  | - |
| Prominent forehead |  |  | + |  | - |
| Deep-set eyes |  |  | + |  | - |
| Short palpebral fissures |  |  | + |  | - |
| Upslanted palpebral fissures |  |  |  |  | 1/3 |
| Epicanthus |  |  |  |  | 2/3 |
| Wide-spaced eyes |  |  |  |  | 1/3 |
| Full cheeks |  |  |  |  | 1/3 |
| Short/anteverted nares |  |  | + |  | - |
| Broad nasal bridge |  |  |  |  | 1/3 |
| Broad nasal tip |  |  | + |  | - |
| Long philtrum |  |  | + |  | - |
| Thin upper lip |  |  | + |  | - |
| Low-set, posteriorly rotated ears |  |  | + |  | - |
| Micrognathia |  |  |  |  | 1/3 |
| <u>Central nervous system findings</u> |  |  |  |  |  |
| Hypotonia | + |  |  |  | 2/3 |
| Muscle weakness/atrophy | + | + |  | + | 1/3 |
| Ataxia | + |  |  |  | 1/3 |
| Seizures | + |  |  | + | - |
| Dyskinesia | + |  |  |  | - |
| Dystonia | + |  |  |  | - |
| Spastic paraplegia | + | + |  |  | - |
| <u>Magnetic resonance imaging findings</u> |  |  |  |  |  |
| Cerebellar hypoplasia/atrophy | + | + | + | + | 1/3 |
| Delayed/abnormal myelination | + | + | + | + | 1/3 |
| Small brainstem/pons | + |  |  |  | 1/3 |
| Enlarged ventricles |  |  | + |  | 2/3 |
| Intracerebellar cysts | + |  |  |  | - |
| Cortical atrophy |  | + |  | + | - |
| Thin corpus callosum |  | + | + |  | - |
| Calcification basal ganglia/thalamus |  |  | + |  | - |
| Enlarged extra-axial spaces |  |  | + |  | 1/3 |
| <u>Ocular findings</u> |  |  |  |  |  |
| Myopia/astigmatism |  |  | + |  | 1/3 |
| Retinitis pigmentosa/retinal dystrophy |  |  | + |  | 1/3 |
| Esotropia |  |  | + | + | 2/3 |
| Corneal dystrophy |  |  | + |  | - |
| Glaucoma |  |  | + |  | - |
| Nystagmus | + |  | + | + | - |
| Persistent pupillary membrane | - |  | - |  | 1/3 |
| Optic atrophy | + |  | - |  | - |
| <u>Cardiac findings</u> |  |  |  |  |  |
| Complete heart block | - |  |  |  | 1/3 |
| <u>Skeletal findings</u> |  |  |  |  |  |
| Craniosynostosis |  |  |  |  | 1/3 |
| Pectus carinatum |  |  |  |  | 1/3 |
| Broad thumbs/phalanges |  |  | + |  | - |
| Brachydactyly |  |  | + |  | 1/3 |
| Clinodactyly 5<sup>th</sup> digit |  |  |  |  | 1/3 |
| Fractures/arthrogryposis |  |  |  | + | - |
| <u>Other</u> |  |  |  |  |  |
| Feeding difficulties | + |  |  |  | 2/3 |
| Respiratory insufficiency | + | + |  | + | 1/3 |
| Sensorineural hearing loss |  |  | + |  | - |
| Hypothyroidism |  |  | + |  | - |

In contrast to *EXOSC3* and *EXOSC8* variants, biallelic variants in *EXOSC2*, encoding a cap exosome subunit, result in a less severe syndrome that is rare (only two European families ascertained) comprising mild intellectual disability, severe myopia, early onset retinitis pigmentosa, progressive sensorineural hearing loss, hypothyroidism, short stature, brachydactyly and broadening of the thumbs and terminal phalanges (30). In addition to severe myopia and retinitis pigmentosa, the syndrome linked to *EXOSC2* variants exhibited several vision/eye-related disorders, including nystagmus, strabismus, corneal dystrophy and glaucoma (30). The syndrome linked to *EXOSC2* variants also showed dysmorphic features, with premature aging, sparse/fine hair, a high/prominent forehead, deep-set eyes, short palpebral fissures, a short nose with a concave nasal ridge/anteverted nares, wide nasal base with a broad nasal tip/wide columella, a long philtrum, thin upper lip and low-set/posteriorly rotated ears (30). Brain MRIs showed mildly enlarged extra-axial spaces and mild cortical and cerebellar hypoplasia/atrophy, diffuse dysmyelination and bilateral calcifications in the basal ganglia and thalamus (30).

Finally, biallelic variants in *EXOSC9*, which encodes a ring exosome subunit, are associated with early-onset weakness and respiratory impairment with a progressive, SMA-like motor neuronopathy and cerebellar atrophy (31, 32). One severely affected individual suffered congenital fractures of the long bones and arthrogryposis at birth and developed respiratory failure resulting in demise at 15 months of age (32). Other patients showed milder phenotypes, with congenital esotropia and nystagmus and poor head control that progressed to developmental delays, muscle weakness, oculomotor dysfunction, coordination difficulties and lower motor neuron symptoms (32). MRI findings included cerebellar and cortical atrophy and possible delayed myelination (32). Thus, mutations that impact multiple structural subunits (EXOSC2/3/8/9) of the RNA exosome cause diverse clinical presentations, with a high prevalence of pontocerebellar hypoplasia and SMA-like phenotypes.

The pathogenic mutations in genes that encode structural subunits of the RNA exosome often cause specific amino acid changes that occur in evolutionarily conserved residues (18). A variety of approaches in genetic model organisms have been employed to begin to understand the requirement for these EXOSC proteins in brain development as well as how pathogenic missense mutations alter the function of the RNA exosome. Studies in zebrafish have employed morpholinos or CRISPR (clustered regularly interspaced short palindromic repeats)/Cas9 injections to knock down or remove expression of the orthologous zebrafish exosome subunits (19, 29, 32, 33). One study reported functional rescue of *exosc3* morpholino knockdown with wild-type *exosc3*, while the *exosc3* variants encoding the amino acid changes found in disease showed no rescue (19). As a complement to these knockdown studies in zebrafish, experiments in the budding yeast, *Saccharomyces cerevisiae*, have been employed to model the pathogenic missense mutations that occur in *EXOSC3* and study the functional consequences of the resulting amino acid substitutions (34, 35). In these studies, the amino acid substitutions that occur in EXOSC3 in association with PCH1b were generated in Rrp40, the budding yeast ortholog of EXOSC3, and examined *in vivo* (34, 35). These studies demonstrated that a budding yeast model could be used to assess how amino acid changes in an exosome subunit linked to disease could alter RNA exosome function and provided some of the first evidence that pathogenic mutations that occur in the exosomopathies impact the function of the RNA exosome.

We now present three patients with biallelic variants in *EXOSC5*, encoding a ring exosome subunit, who demonstrate symptom overlap with disease phenotypes characterized in patients with deleterious variants in other RNA exosome subunit genes. Studies in zebrafish support a critical role for EXOSC5 in brain development. We then model the pathogenic amino acid substitutions in the budding yeast EXOSC5 ortholog, Rrp46, as well as mammalian EXOSC5. None of the mammalian EXOSC5 proteins show significantly altered protein levels, suggesting distinct molecular mechanisms that impact the function of the essential RNA exosome. Consistent with this conclusion, distinct patterns of RNA processing defects are evident in the yeast model. Studies in cultured neuronal cells show differences in interactions between EXOSC5 harboring pathogenic amino acid substitutions and other RNA exosome subunits. This work adds an additional structural subunit of the RNA exosome to the expanding list of exosome subunit genes for which mutations have been linked to human disease, while also providing evidence for distinct consequences of the different pathogenic *EXOSC5* variants.

## Results

### Clinical Presentations

#### Patient 1

The first patient identified (Fig. 1) was a 10-year old female. Both parents were healthy and there was no relevant family history, although her father was reportedly treated with medication for obsessive-compulsive disorder. Paternal ethnicity was Cherokee, Mexican and French and maternal ethnicity was English, Portuguese, Native American and Spanish. There was no known consanguinity. At the time of the birth, the mother was 22 years of age and a primigravida and the father was 23 years of age. The pregnancy was uneventful, but resuscitation was needed after a difficult delivery due to a nuchal cord. At 4 months of age, the patient had no head control and was unable to roll. She was unable to crawl at 11 months of age and was diagnosed with developmental delays and ataxia. An MRI of the brain showed cerebellar hypoplasia, with a small cerebellar vermis, enlarged cerebellar sulci and a small brainstem (Fig. 1C). At 2 years of age, investigations for cyanotic episodes and syncope showed a heart rate of 30 beats per minute with complete heart block and pacemaker insertion was performed. There were no structural cardiac defects on echocardiogram. At 3 and a half years of age, she used head movements rather than eye movements to look for objects. An ophthalmological examination showed left esotropia, a right alternating esotropia, myopia and astigmatism. Examination of the fundus showed thin vessels, diffuse changes in the retinal pigment epithelium at the periphery and optic disc pallor (Fig. S1). Optical coherence tomography revealed attenuation of the outer retinal layers that was consistent with a generalized retinal degeneration (Fig. S2). An electroretinogram (ERG) recorded under general anesthesia using Burian-Allen contact lens electrodes showed diffuse cone greater than rod dysfunction in each eye (Fig. S3A-D) and she was treated with visual aids and surgery for the esotropia. The ERG was repeated at 6 years of age and showed progressive reduction of cone responses with similar rod responses, consistent with progressive cone-rod dystrophy (Fig. S3E-H). Visual acuity was measured at 20/125 in each eye at 9 years of age whilst wearing glasses (−4.00+3.25×090 right eye, -3.75+2.00×090 left eye). At 10 years of age, she attended a special education class with an individualized education plan and received physical therapy, occupational therapy and speech therapy. She had relatively mild speech delays and could communicate in short phrases and sentences and had good comprehension. She had developed an obsessive-compulsive disorder, with an emphasis on cleanliness and routines, that was treated with medication. She also required medication for difficulty with sleep, but a sleep study was normal.

**Figure 1.**
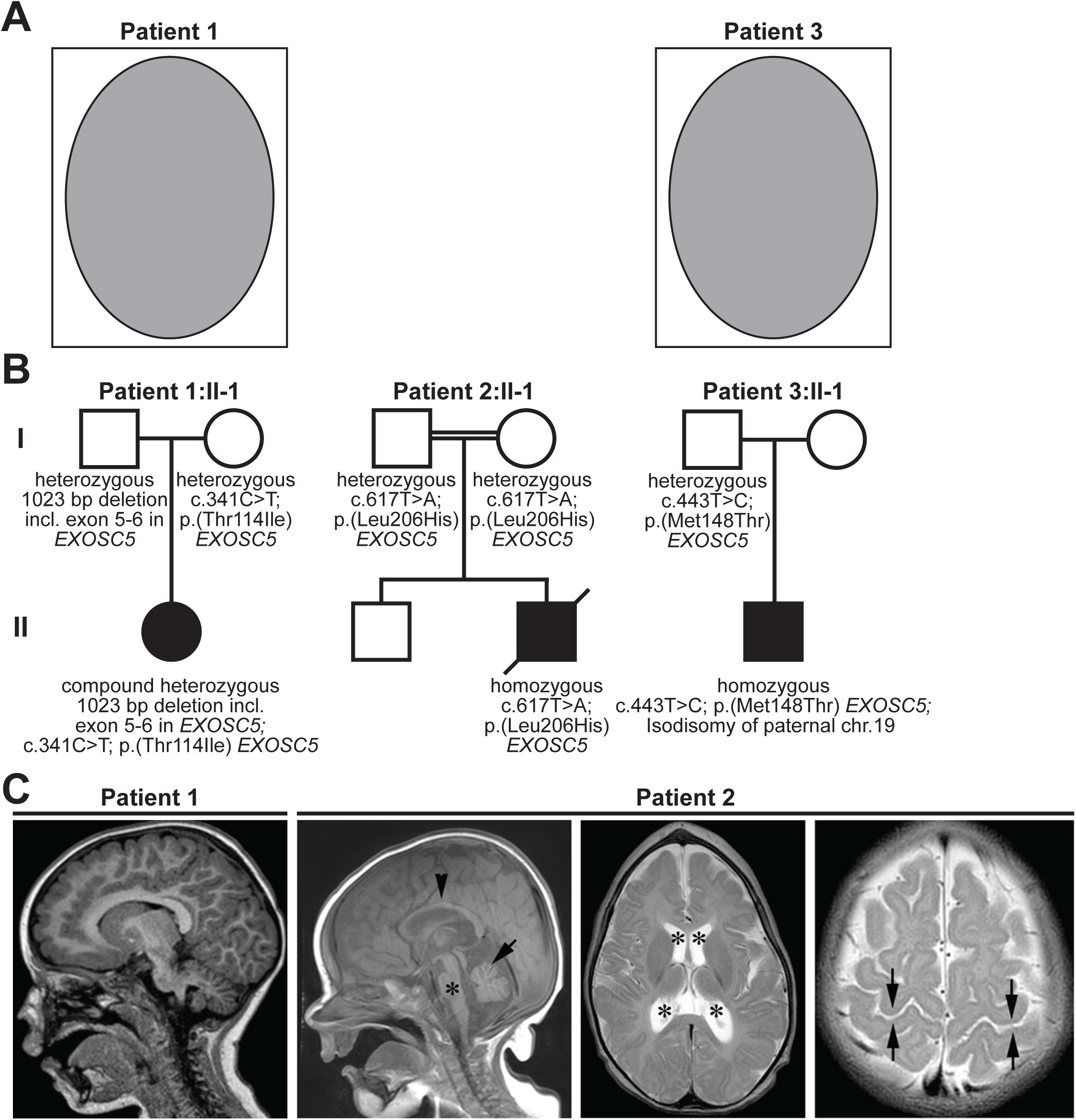
Clinical presentation and pedigrees of three patients with biallelic variants in *EXOSC5*. (A) (redacted) Facial photographs of Patient 1, demonstrating mild ptosis, epicanthic folds, full cheeks, a high nasal bridge, broad nasal tip, prominent Cupid’s bow and mild micrognathia, and Patient 3, demonstrating a mildly broad nasal tip and prominent Cupid’s bow. Patient 3 was not considered to have significant facial anomalies by the Attending clinician. (B) Pedigrees illustrating the inheritance pattern of *EXOSC5* alleles in Patients 1, 2, and 3 are shown. (C) Magnetic resonance imaging (MRI) findings in *EXOSC5* Patients 1 and 2 are shown. MRI scan of the brain of Patient 1 with p. Thr114Ile, showing cerebellar hypoplasia, with reduced size of the cerebellar vermis and increased cerebellar sulci on sagittal section. The brainstem was also small. MRI scan of the brain of Patient 2 with homozygosity for an *EXOSC5* variant p. Leu206His at the age of 4 months. Left panel: Sagittal T1-weighted MRI revealed decreased size of pons and brainstem (asterisk) and also of the cerebellar vermis (arrow). The posterior fossa was enlarged. The corpus callosum (arrowhead) appeared thin, but was within the expected physiological variation for the age. Middle panel: Axial T2-weighted MRI illustrated the coned shape of the skull and the corresponding coned shape of both frontal lobes, due to early closure of the metopic suture. The size of the ventricular system is increased (black stars). Right panel: Axial T2-weighted MRI illustrated the relative lack of reduced T2-signal intensity in the perirolandic areas (black arrows), indicating hypomyelination.

The patient was last examined at 10 years and 10 months of age. Height was 128.5 cm (2^nd^ centile), weight was 31.8 kg (24^th^ centile) and occipitofrontal circumference (OFC) was 52.8 cm (58^th^ centile). She had minor anomalies that included mild ptosis, small epicanthic folds, a high nasal bridge with a broad nasal tip, full cheeks, mild retrognathia, and fifth finger brachydactyly and clinodactyly. No nystagmus was observed. She was able to stand holding on and balance without support for several seconds. She could take 3-4 steps alone, but was unsteady and had an ataxic gait that necessitated a walker or a wheelchair for ambulation. Her speech was slurred and she had past pointing when she reached for objects. Prior genetic testing included a negative microarray (Agilent platform with 105,072 60-mer oligonucleotide probes with a spatial resolution of 10-35 kb) and normal carbohydrate deficient glycoprotein testing. Sequencing for 58 confirmed, disease-associated mitochondrial DNA mutations and deletions revealed no reportable variants (GeneDx, Inc.).

#### Patient 2

Patient 2 was a male and was the second child born to first cousins from Iraq (Fig. 1). At the time of birth, the mother was 30-years of age and the father was 36-years of age. The patient’s mother had thalassemia minor and the older sibling was diagnosed with phenylketonuria. After an uneventful pregnancy, the patient was born at 41+3 weeks of gestational age by breech presentation. Birth measurements were weight 2,930 g (5^th^ centile), OFC 34.5 cm (28^th^ centile) and crown-rump length 35 cm (mean to +2 standard deviations). At one week of age, length was 48 cm (4^th^ centile). He was noted to have trigonocephaly, upslanted palpebral fissures, ophthalmoplegia, esotropia, widely-spaced eyes, epicanthic folds, a broad nasal root, micrognathia, pectus carinatum, full cheeks and accessory mamillae. In the neonatal period, he was fed with a nasogastric tube and breathing was supported with a continuous positive airway pressure (C-PAP) apparatus. At 2 months of age, the patient required gastrostomy and tracheostomy. Growth parameters were within the normal ranges.

At the time of birth, he was able to move his upper arms against gravity, was able to move his legs and had some head control, but his spontaneous movements were reduced. Soon after birth, moderate hypotonia, including medialized vocal chords, was diagnosed. He had reduced grasp and Moro reflexes and the sucking reflex was likely absent. His spontaneous movements decreased during infancy, indicating regression, and at 11 months of age, he was only able to move his fingers. However, he had preserved facial expression and was able to open his eyes and cry. He responded to physical contact and sound stimuli with short moments of smiling. His clinical course was complicated by recurrent airway infections and progressive respiratory impairment. He died at 11 months of age due to a viral infection leading to respiratory failure, gastric retention and reduced bowel function.

Neurophysiological examination performed at 1 month of age revealed subacute axonal motor neuropathy, with ongoing denervation in both proximal and distal muscles in upper and lower extremities, resembling SMA. Sensory responses were normal. A histological examination of a biopsy from the vastus lateralis muscle showed neurogenic and myopathic changes, with fiber size variation consisting of hypertrophic and atrophic/hypotrophic round fibers of both types, but mainly type 2 fibers. A brain MRI at the age of 4 months showed trigonocephaly with premature fusion of the metopic sutures, reduced size of cerebellar vermis, brainstem and pons, enlargement of the ventricular system, and hypomyelination; a thinned corpus callosum was also noted, but was considered within the physiological range for age (Fig. 1C). An MRI scan of the spinal cord was normal. Electroencephalograms (EEGs) performed at 1 and 4 months of age did not identify definite abnormalities.

At 2 months of age, flash-visual evoked potential (VEP) testing was normal. Ophthalmologic examination at 9 months of age showed a normal fundus with unilateral persistent pupillary membrane and bilateral closure deficit. Esotropia and latent nystagmus were noticed. He had uncontrolled eye movements, and the degree of eye contact and fixation was uncertain. Cardiological examination revealed a normally structured heart with a slightly enlarged right side, and electrocardiography was normal. Metabolic screening of blood did not detect abnormalities and a microarray analysis (1M aCGH Agilent Technologies) did not reveal a clear pathological finding.

#### Patient 3

The third patient was a 17-month old male (Fig. 1). The pregnancy was complicated by pre-eclampsia and an ultrasound scan showed that the length of the long bones measured just below the lower limit of normal at 35 weeks of gestation. The baby was delivered by C-section at 33 4/7 weeks of gestation. The neonatal course was complicated by neonatal anemia, mild respiratory distress, which resolved, and neonatal osteopenia. He was diagnosed with oropharyngeal dysphagia and laryngomalacia, which both later resolved. Early milestones were notable for fine and gross motor delays and physical therapy and occupational therapy were commenced. An MRI scan of the brain at 9 months of age showed increased size of the lateral and third ventricles, with the lateral ventricles measuring 1 cm at the frontal horn and the third ventricle measuring 0.9 cm. Septation of the inferior aspect of the cerebral aqueduct was suspected and there was benign enlargement of the extra axial spaces. A spinal MRI showed that the conus medullaris terminated at L2. An echocardiogram showed an anomalous coronary artery fistula. Additional clinical findings included penoscrotal hypospadias, a tethered cord and a sacral dimple. At 12 months chronological age (10.5 months corrected age), he could commando crawl, cruise and pull to stand with assistance, but was unable to get from lying down to sitting or lower himself to the ground when standing. He was able to rake and feed himself small items, but did not have a pincer grip. He turned to his name, but was not inhibited by ‘no’ and he had no stranger anxiety. His development was assessed as age appropriate for language, cognitive and social skills, but had fine motor delays and his motor developmental age was assessed as consistent with 8 months of age. When last examined at 14 months of age, length was 69.7 cm (0^th^ centile), weight was 9.3 kg (14^th^ centile) and OFC was 47.5 cm (87^th^ centile). He had no dysmorphic features. A single nucleotide polymorphism (SNP) array (Affymetrix human SNP array 6.0) showed paternal isodisomy for chromosome 19, arr[GRCh37] 19p13.3q13.43 (1,402,996-59,097,752)×2 hmz, without any copy number variants. The homozygous region contained an estimated 2,200 genes, of which approximately 110 were thought to be associated with autosomal recessive disorders.

### Identification of *EXOSC 5* Variants

Exome sequencing in Patient 1 showed a paternally inherited, partial gene deletion of 1,023 bp that includes exons 5-6 in *EXOSC5*, reported as arr[GRCh37]19q13.2(41,892,557-41,893,580)x1, that was confirmed by microarray (Table 3; Fig. S4). Deletions of this chromosome region have not previously been described (Database of Genomic Variants, http://dgv.tcag.ca and Decipher, https://decipher.sanger.ac.uk). Patient 1 also had a maternally inherited, missense variant, chr19(GRCh37):g.41897789;NM_020158.3:c.341C>T;p.Thr114Ile, in *EXOSC5* (NM_020158.3; Fig. S5). This variant has been found at low frequencies in control databases (9/11,498 in the ExAC database and 17/232,236 in the gnomAD database, with a maximum allele frequency of 0.06% in the Latino population) and is absent from the 1000 Genomes database (Table 3). Database predictions favored pathogenicity, with a combined annotation dependent depletion (CADD) score of 27 (36) and designations of probably damaging (p = 1.0) and disease-causing (p = 0.999) from PolyPhen2 (37) and MutationTaster (38), respectively (Table 3). The threonine and isoleucine residues differ in polarity, charge and size and the threonine residue is conserved in diverse animals, fungi and plants (Fig. S6; >85% of the sequences contain Thr or Ser). Patient 1 also had a maternally inherited, heterozygous variant in the ATP-binding cassette, subfamily A, member 4 (*ABCA4*) gene, c.2588G>C, predicting p.Gly863Ala, that was reported as pathogenic by the testing laboratory, but a second variant in *ABCA4* was not identified. Her mother had no ocular symptoms. This variant has a population frequency of 1214/282648, or 4.3 × 10^−3^ and has been reported in the homozygous state 7 times in gnomAD. On the basis of this high allele frequency and the lack of symptoms in her mother, we considered that this variant was unlikely to be responsible for the proband’s cone-rod dystrophy.

The second patient was homozygous for chr19(GRCh37):g.41892629A>T;NM_020158.3:c.617T>A; p. Leu206His in *EXOSC5* (Table 3; Fig. S5). Database predictions regarding the pathogenicity of this variant were suggestive of a deleterious effect, with a CADD score of 29.3, SIFT score of 0 and MutationTaster designation of disease-causing, but the PolyPhen2 score was 0.242, consistent with a benign variant (Table 3). This variant was absent from control databases, including gnomAD, and the residue was conserved in diverse animals, fungi and plants (Fig. S6; all of the sequences contain Leu, Phe, Tyr or Trp).

The third patient had uniparental isodisomy for the paternal chromosome 19 and was homozygous for a paternally inherited, missense variant, chr19(GRCh37):g.41895752A>G;NM_020158.3:c.443T>C;p. Met148Thr in *EXOSC5* (Table 3). This variant was predicted to be pathogenic by Polyphen2, MutationTaster and CADD and was not found in homozygous form in any public database (Table 3).

*EXOSC5* has a probability of loss of function intolerance (pLI) score of 0, with 9.2 predicted loss of function variants and 5 observed (gnomAD). However, no homozygous variants have been reported in the exons of this gene in control databases and there are no entries for pathogenic or likely pathogenic variants in ClinVar.

### CRISPR/Cas9 Model of *exosc5* Loss of Function in Zebrafish

To explore the requirement for *EXOSC5* in brain development, we employed zebrafish as previously used for models of *EXOSC3, EXOSC8* and *EXOSC9* loss of function (19, 29, 32, 33). The zebrafish orthologous gene, *exosc5*, has two predicted transcripts – the longest transcript has 6 exons and predicts a 222 amino acid protein, (ENSDART00000059352.6), whereas the shorter one is predicted to be 128 amino acids in length (ENSDART00000145177.2). The zebrafish *exosc5* cDNA sequence shows 58% identity with the 235 amino acid human gene (NM_020158.4) (Fig. S6) and has a gene order conservation score of 50 (Ensembl). This conservation supports the use of model systems to explore the function of *EXOSC5*, with zebrafish employed to define the requirement for *exosc5* in neurodevelopment.

Larvae injected with sgRNA/Cas9 targeting exon 2 of *exosc5* (Fig. S7) show increased tail curvature, shortening of the tail and body, edema, small eyes and heads and fin defects compared to EKW controls (Fig. 2A). We classified larvae into three categories of severity based on larval phenotypes reported from the previous study targeting *exosc9* with antisense morpholinos and CRISPR/Cas9 (32). Larvae with mild tail curvature were designated as ‘mild phenotype’, those with hooked tail curvature were designated as ‘moderate phenotype’ and larvae with severe shortening of the tail and body axis were designated as ‘severe phenotype’ (Fig. 2B). To confirm the loss of exosc5, we cloned and sequenced PCR products of the *exosc5* target locus from multiple larvae with mosaicism. This analysis revealed multiple indel variant alleles of *exosc5* from a single PCR product and confirmed the absence of any wild-type *exosc5* allele (Fig. S7), consistent with biallelic inheritance and predicted loss of *exosc5* function. As shown in Figure 2C, representative larvae with biallelic indel variants in *exosc5* demonstrated reduced size of the eyes, head and brain mesencephalon and telencephalon.

**Figure 2.**
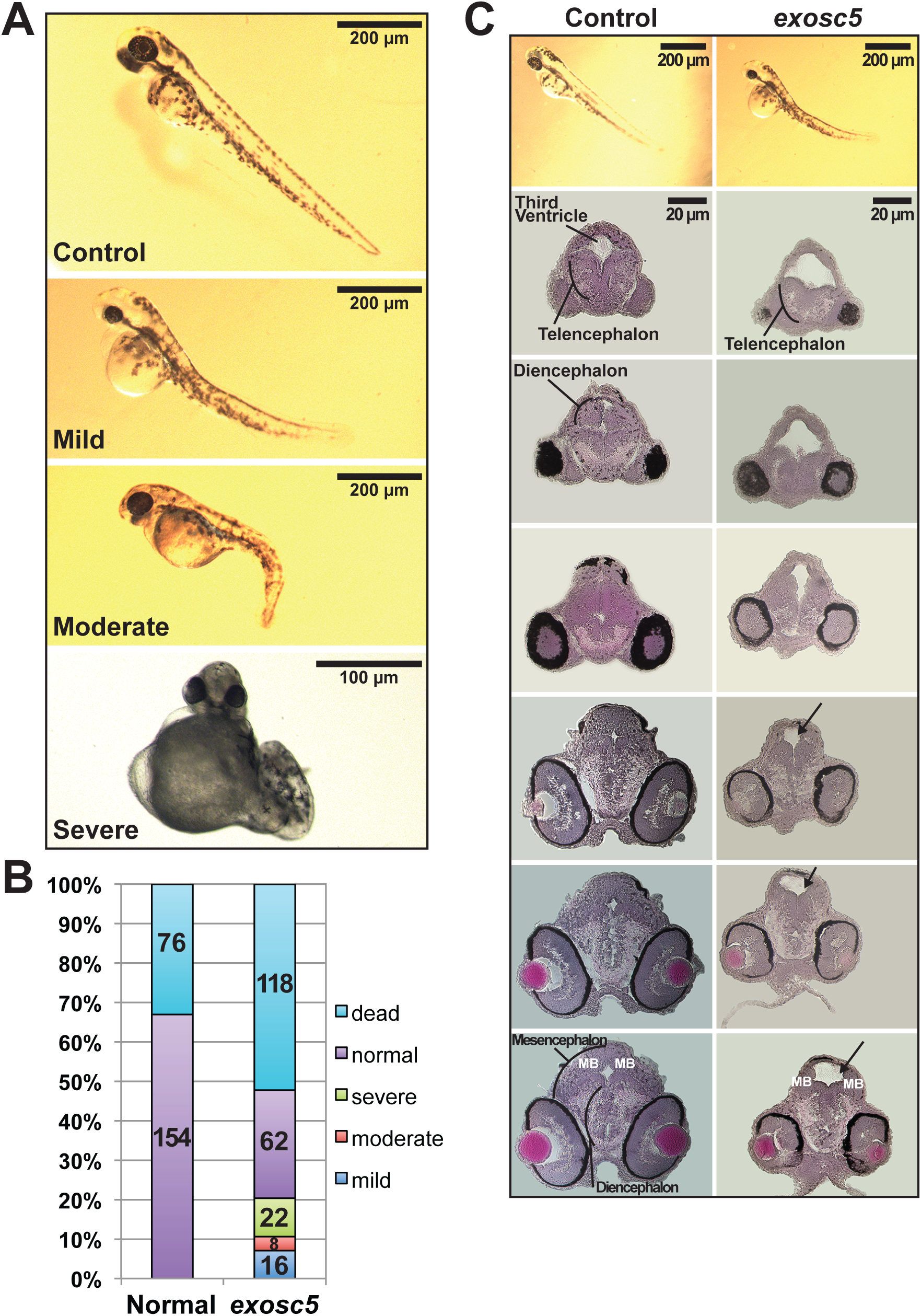
CRISPR/Cas9 targeting of *exosc5* in transgenic zebrafish demonstrates larvae with morphological defects of the tail and body. (A) Typical morphology was observed in Control larvae. For CRISPR-injected larvae with targeting of *exosc5*, we observed larvae with a mild tail curvature (Mild phenotype), hooked tail curvature (Moderate phenotype); and larvae with severe shortening of the tail and body axis with distortion of the body axis (Severe phenotype). Reduced eye and head size and edema were also observed in the *exosc5*-targeted larvae. Larvae were photographed at 54-58 hours post fertilization (hpf). (B) The relative number of morphological phenotypes (Normal, Dead, Mild, Moderate, and Severe) observed in Control (n=230) and *exosc5* crispants (n=226) is depicted as percentage of total larvae analyzed. (C) Sections of Control larvae and CRISPR-injected larvae with targeting of *exosc5* were stained with hematoxylin and eosin. Crispant *exosc5* larvae showed reduced eye and lens size, increased ventricle size, and reduced brain size with aberrant formation of the mesencephalon and diencephalon as indicated. Larvae were photographed at 54-58 hpf.

As a complement to examining morphology, we employed an *mnx1* transgenic strain driving enhanced green fluorescent protein (EGFP) expression in motor neurons (39) and an NBT:DsRed strain that labels neurons under the *neural ß-tubulin* promoter (40) (Fig. 3). CRISPR injections targeting *exosc5* in *mnx1* and NBT zebrafish strains yielded similar results to injections in control EKW zebrafish, with an overrepresentation of mild, moderate and severe phenotypes compared to controls (data not shown). In the *mnx1:EGFP* strain with EGFP labeling of motor neuron synapses, *exosc5* targeting causes severe phenotypes as illustrated by reduced fluorescence in the somatic neurons of the midbrain and hindbrain (Fig. 3A). In the NBT strain with a pan-neuronal marker, *exosc5* targeting causes variable brain morphology in larvae with severe phenotypes, with relative preservation of brain and spinal cord morphology in one larva compared to complete disruption of brain morphology in another (Fig. 3B). qRT-PCR experiments performed on RNA obtained from abnormal larvae in *exosc5*-targeted CRISPR injections showed *exosc5* expression was significantly reduced in the CRISPR-injected larvae compared to controls (Fig. S8). These studies reveal a critical role for *EXOSC5* in neurodevelopment.

**Figure 3.**
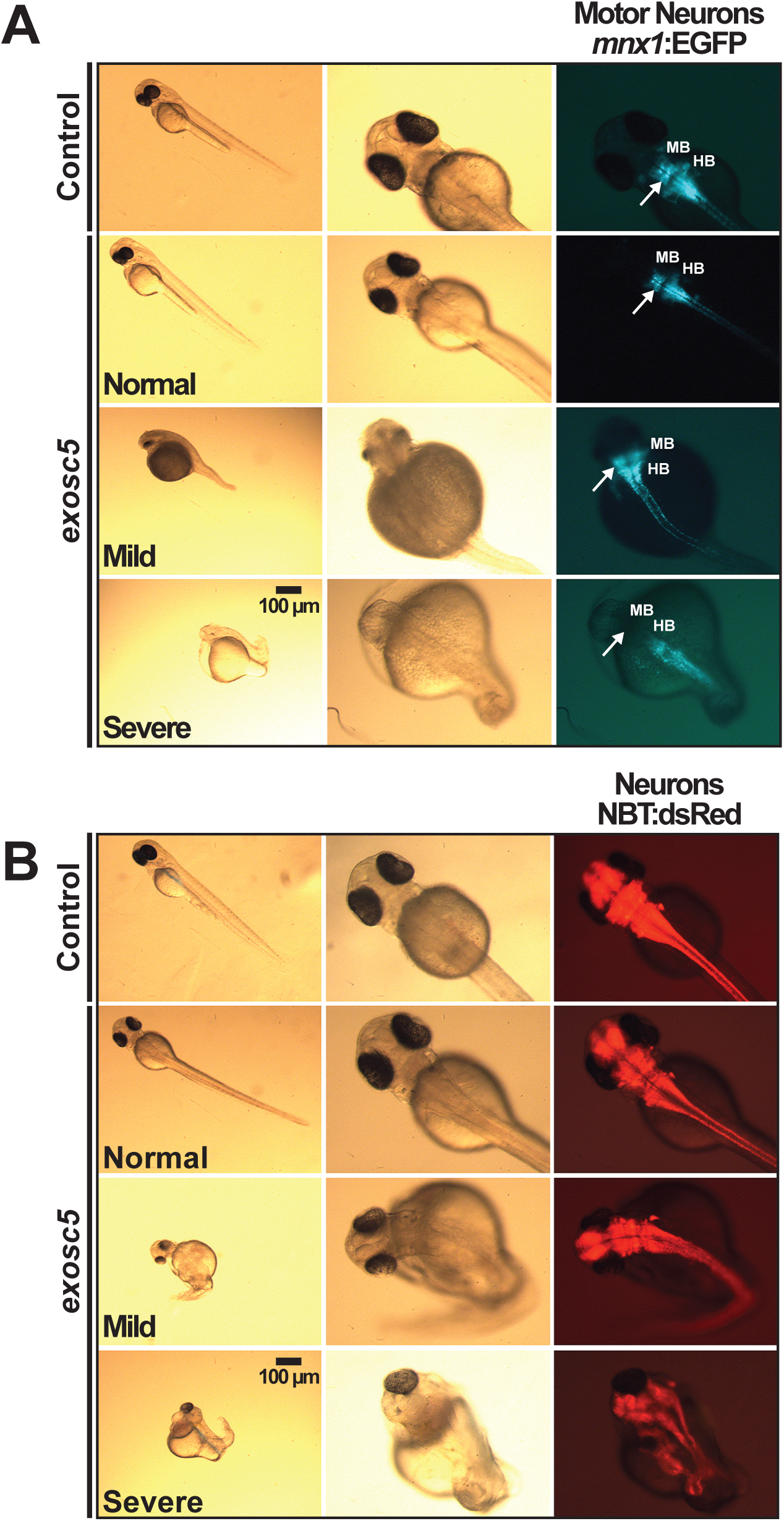
CRISPR/Cas9 targeting of *exosc5* in *mnx1*:EGFP and NBT:dsRed transgenic zebrafish demonstrates larvae with morphological defects of the tail, body and brain. (A) Transgenic zebrafish with *mnx1*:EGFP with sgRNA targeting of *exosc5* show severe larval phenotypes, with shortened heads, bodies and tails, small or absent eyes and edema compared to Control. Larvae with targeting of *exosc5* can also have a normal appearance similar to controls as illustrated in the Normal panel. Fluorescence imaging reveals decreased GFP signal and abnormal midbrain and hindbrain morphology in larvae with Severe phenotypes, marked by arrows, compared to Control and normal-appearing injected larvae. Larvae with Severe phenotypes were verified to have biallelic *exosc5* targeting by Sanger sequencing. Larvae were photographed at 54-58 hpf. MB = midbrain; HB = hindbrain. (B) Transgenic zebrafish with Neurons labeled by NBT1:dsRed with sgRNA targeting of *exosc5* show severe larval phenotypes, with shortened heads, bodies and tails, distorted bodies, small eyes and marked edema compared to Control. Larvae with targeting of *exosc5* can also have a Normal or Mild phenotype. Fluorescence imaging reveals extensive disruption of brain formation and morphology in one *exosc5*-targeted larvae, which shows a Severe phenotype. Larvae with severe phenotypes were verified to have successful *exosc5* targeting by Sanger sequencing. Larvae were photographed at 54-58 hpf.

### Functional consequences of pathogenic amino acid changes in EXOSC 5

All patients identified are either homozygous (Patients 2 and 3) or heterozygous (Patient 1) for missense variants that encode single amino acid substitutions in the EXOSC5 protein (Fig. 4), raising the question of how these changes impact RNA exosome function. The EXOSC5 protein is a structural subunit of the 10-subunit RNA exosome complex and one of the six subunits that compose the core ring of the complex (Fig. 4A). EXOSC5 and the other RNA exosome core ring subunits primarily contain a single RNase PH-like domain that resembles the bacterial enzyme RNase PH (phosphate nuclease), but lacks catalytic activity (7) (Fig. 4B). As illustrated in Figure 4B, the amino acid changes present in EXOSC5 in patients occur in regions of the protein that are evolutionarily conserved between humans, zebrafish and budding yeast, although only Leu206 is strictly conserved as Leu191 in budding yeast. To begin to explore how these amino acid substitutions could alter the structure of EXOSC5 or the contacts that EXOSC5 makes with other RNA exosome subunits within the core complex, we took advantage of the RNA exosome structural model (41). Each of the amino acids altered is located in a different region of the EXOSC5 protein (Fig. 4C (Panel II)). Two of these residues, Thr114 and Leu206, are located in regions of EXOSC5 that contact other RNA exosome subunits (Fig. 4C (Panel I)). As shown in Figure 4C Panel IV, the p.Thr114Ile amino acid substitution could result in the loss of a hydrogen bond that the EXOSC5 Thr114 residue makes with the backbone of the EXOSC5 Ala62 residue. This is a minor change that is not predicted to have a significant effect on the stability of the EXOSC5 chain. Instead, most likely, the effect of the p.Thr114Ile substitution can be attributed to the central location of this residue within the RNA exosome complex and proximity to EXOSC3 (Fig. 4C, panel IV), as the mCSM predicted a -0.42Kcal/mol (destabilizing) change in the affinity of the protein-protein interaction with EXOSC3. The p.Leu206His amino acid change replaces a buried hydrophobic residue with a polar residue (Fig. 4C, Panel III). This change is predicted to reduce EXOSC5 protein stability (−1.0 Kcal/mol according to mCSM) and exert a destabilizing effect on the protein-protein interaction with EXOSC8 (−0.7 Kcal/mol according to mCSM). The p.Met148Thr substitution has an even more dramatic predicted effect on protein stability (−2.2 Kcal/mol according to mCSM) and interaction with EXOSC3 (−1.1 Kcal/mol according to mCSM). This structural analysis suggests that the amino acid changes present in EXOSC5 found in the three patients have the potential to alter the stability or conformation of the EXOSC5 protein, possibly affecting interactions with neighboring subunits.

**Figure 4.**
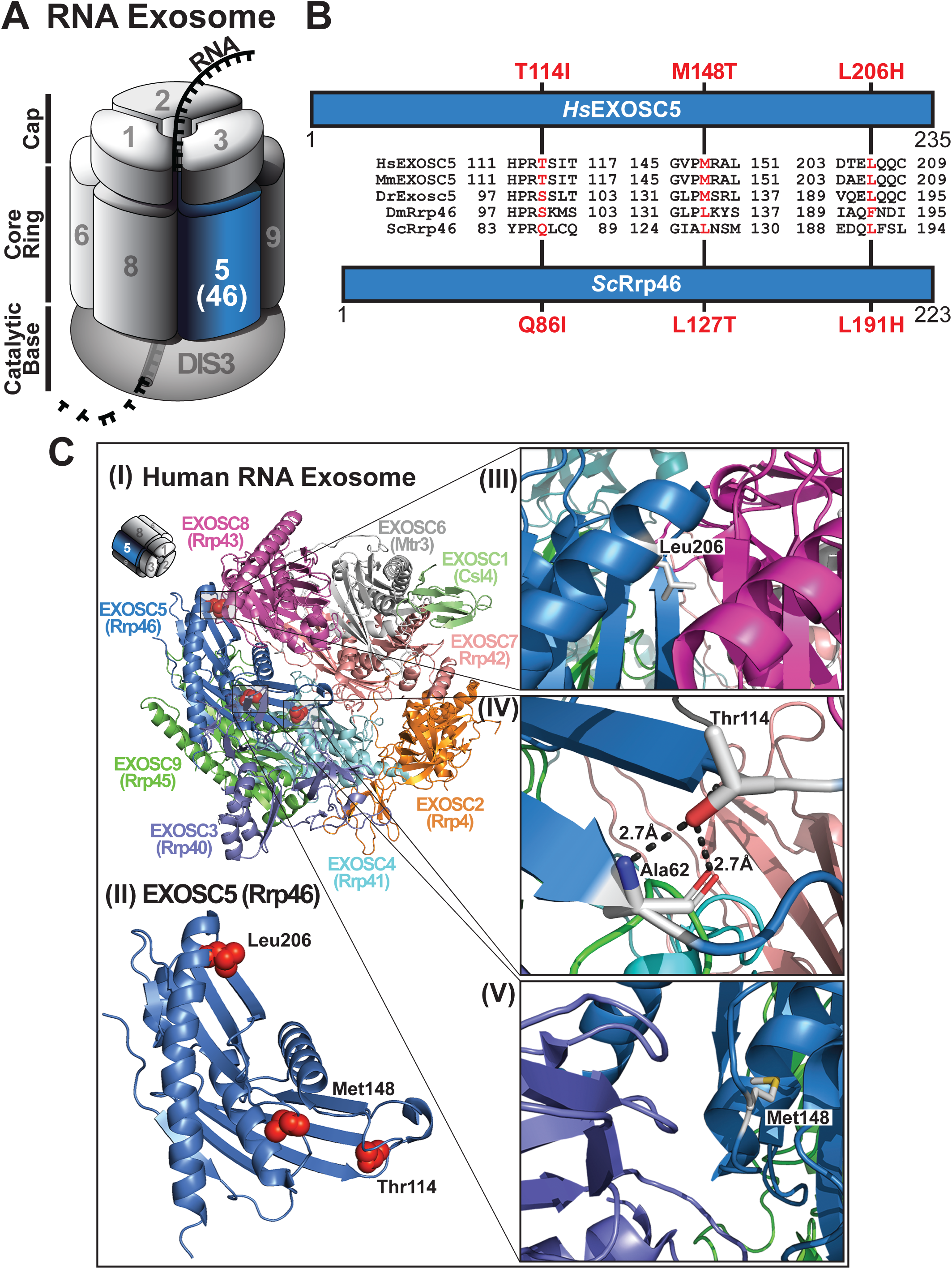
The amino acid changes in EXOSC5 identified in patients lie in conserved regions of the protein, which could be important for interactions within the RNA exosome complex. (A) A cartoon model of the 10-subunit RNA exosome complex is shown. The subunits of the complex are indicated by their EXOSC number (1-3 in the cap, 4-9 in the core ring), with the catalytic DIS3 subunit at the base. EXOSC5, which is a component of the core ring structure is highlighted in marine blue. (B) Domain structure schematics are shown for both human EXOSC5 and the *S. cerevisiae* ortholog of EXOSC5, Rrp46. Both proteins are comprised of a single RNase PH-like domain. The amino acids changes - T114I, M148T, and L206H - in EXOSC5 identified in the patients described here are indicated above EXOSC5 and the corresponding amino acid changes - Q86I, M127T, and L191H - in Rrp46 are denoted below Rrp46. Between the domain structures, alignments of the amino acid sequences from human (*Hs*), mouse (*Mm*), zebrafish (*Dr*), fruit fly (*Dm*), and budding yeast (*Sc*) EXOSC5/Rrp46 orthologs that surround the EXOSC5 residue altered in patients (indicated in red) are shown. The alignments highlight the evolutionary conservation of the residue altered in disease and the residues adjacent to it. (C) Structural model of (I) the human 9-subunit RNA exosome complex and (II) the EXOSC5 subunit are shown with the position of the residues altered in patients highlighted as red side chain spheres. A miniature cartoon of the RNA exosome in the top left corner denotes the approximate orientation of the RNA exosome structure. The cap subunits EXOSC1/2/3 are depicted in lime/orange/slate blue and the core ring subunits EXOSC4/5/6/7/8/9 are depicted in cyan/marine blue/gray/salmon red/magenta/forest green, respectively. The depicted human 9-subunit RNA exosome structure is adapted from the human 10-subunit RNA exosome-MPP6 complex (PDB: 6h25)(41) and does not show DIS3 or MPP6. A zoomed-in view of the location of the (III) EXOSC5 Leu206 residue that is changed to Histidine in patient 2 (p.Leu206His) shows that Leu206 is a buried hydrophobic residue and predicts that its replacement with a polar Histidine residue would alter EXOSC5 stability and exert a destabilizing effect on its interaction with EXOSC3. A zoomed-in view of the location of the (IV) EXOSC5 Thr114 residue that is changed to Isoleucine in patient 1 (p.Thr114Ile) shows that Thr114 makes a hydrogen bond with the backbone of the Ala62 residue in EXOSC5 and predicts that its replacement with Isoleucine would disrupt this interaction with Ala62. However, the Isoleucine residue at position 114 in EXOSC5 is not predicted to significantly affect protein stability, but more likely exerts a deleterious effect due to its central location in the exosome and proximity to EXOSC3. A zoomed-in view of the location of the (V) EXOSC5 Met148 residue that is changed to Threonine in patient 3 (p.Met148Thr) predicts that Met148 substitution with a polar Threonine residue would reduce the stability of EXOSC5 and affect its interaction with EXOSC3 (Thr81).

To investigate the functional consequences of the T114I, M148T, and L206H amino acid substitutions in EXOSC5, we engineered the corresponding Q86I, L127T and L191H amino acid changes in the budding yeast ortholog of EXOSC5, Rrp46 (6) (Fig. 4B). We expressed wild-type Rrp46 and variants rrp46-Q86I, rrp46-L127T, rrp46-L191H in budding yeast cells as the sole form of the essential Rrp46 protein, allowing us to assess how these amino acid substitutions impact the function of Rrp46. First, we tested whether each rrp46 variant can support yeast cell growth using a serial dilution and spotting assay. As shown in Figure 5A, *rrp46-Q86I* and *rrp46-L127T* mutant cells show growth similar to control *RRP46* cells at all temperatures tested. In contrast, *rrp46-L191H* mutant cells show detectable growth defects at 30°C that are more severe at 37°C (Fig. 5A). We complemented these solid media growth assays with growth curves in liquid media (Fig. 5B), which confirm that *rrp46-L191H* cells have significant growth defects at 30°C and 37°C compared to the control *RRP46* cells or mutant *rrp46-Q86I* and *rrp46-L127T* cells. Indeed, the doubling time for the *rrp46-L191H* cells at 30°C is ∼ 133 minutes, 64% longer than the control, while the doubling time at 37°C is ∼ 169 minutes, 69% longer than the control.

**Figure 5.**
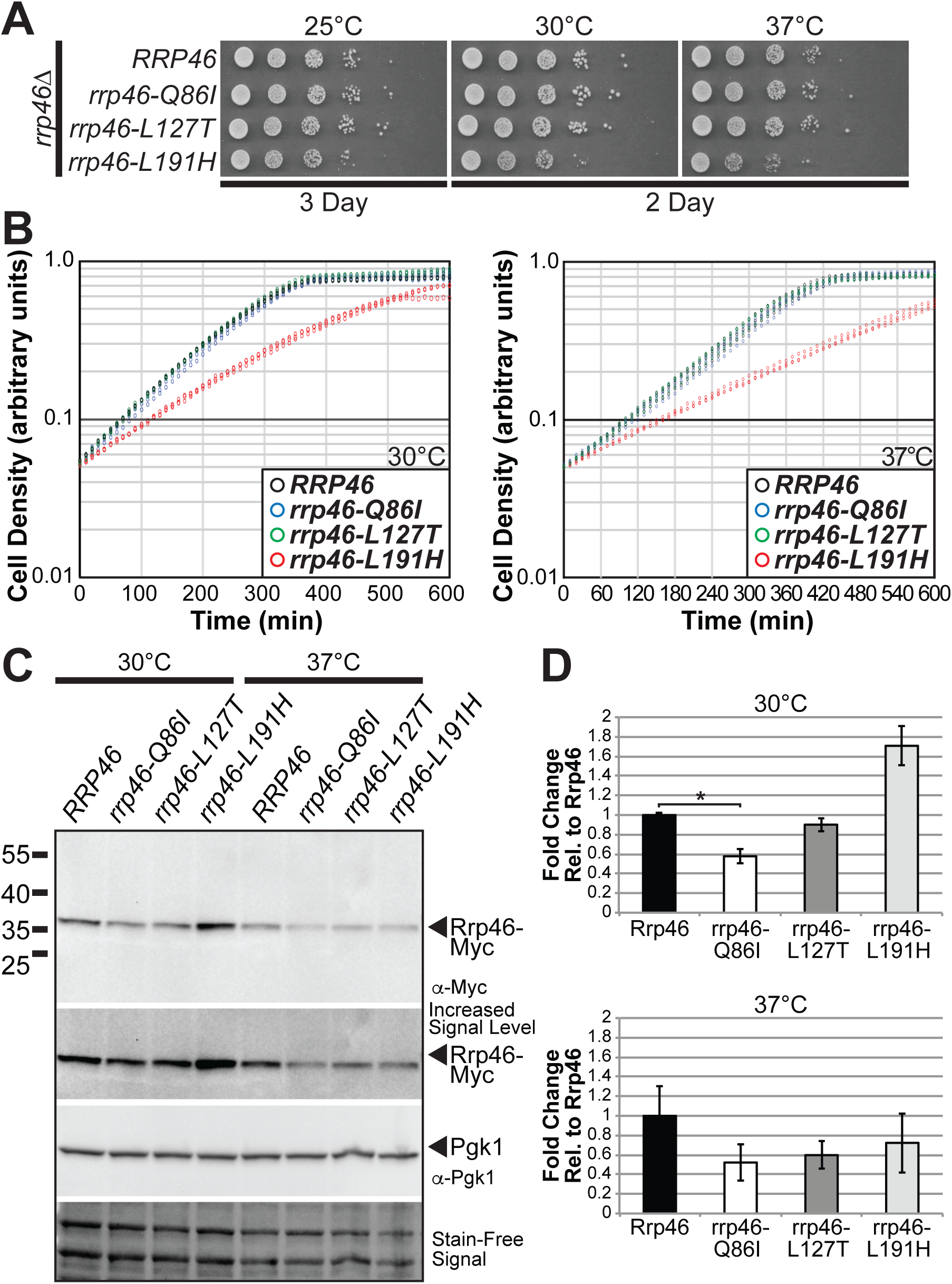
Amino acid substitutions in the budding yeast ortholog of EXOSC5, Rrp46, that correspond to the EXOSC5 T114I, M148T, and L206H substitutions identified in patients impair yeast cell growth. (A) Budding yeast cells that express the rrp46-L191H variant, corresponding to the EXOSC5 p.Leu206His variant, as the sole copy of the essential Rrp46 protein show impaired growth at 30°C and 37°C compared to cells expressing wild-type Rrp46 in a solid media growth assay. In contrast, cells expressing the rrp46-Q86I or rrp46-L127T variant, corresponding to the EXOSC5 p.T114I and p.M148T variant, show no growth defect at any temperature compared to cells expressing wild-type Rrp46. Growth of *rrp46Δ* yeast cells containing *RRP46, rrp46-Q86I, rrp46-L127T*, or *rrp46-L191H* plasmid was analyzed by serial dilution, spotting on solid media, and growth at indicated temperatures. (B) The *rrp46-L191H* mutant cells, but not the *rrp46-Q86I* or *rrp46-L127T* mutant cells, exhibit reduced growth at 30°C and 37°C compared to wild-type *RRP46* cells in a liquid culture growth assay, in which the optical density of the cultures was measured over time. (C) was employed to assess the steady-state levels of the Rrp46/rrp46 proteins. Lysates of *rrp46Δ* yeast cells expressing only Myc-tagged wild-type Rrp46, rrp46-Q86I, rrp46-L127T, or rrp46-L191H grown 30°C or 37°C as indicated were analyzed by immunoblotting with anti-Myc antibody to detect Myc-tagged Rrp46 and rrp46 variants (Rrp46-Myc) and anti-Pgk1 antibody to detect 3-phosphoglycerate kinase (Pgk1) as a loading control. A representative result is shown. (D) Results from two independent immunoblotting experiments (C) were quantitated as described in Materials and Methods. The amount of wild-type Rrp46 protein detected at grown 30°C (top) or 37°C (bottom) was set to1.0 and the normalized fold-change of each rrp46 protein relative to wild-type Rrp46 is shown for each temperature. A statistically significant changes in steady-state protein level was detected for the rrp46-Q86I protein at 30°C (*p* = 0.0307), while the fold change in rrp46-L191H at 30°C was not statistically significant (*p* = 0.0710), and the remaining fold changes in rrp46 levels at 30°C and 37°C were not statistically significant. * indicates *p*-value <0.05.

A growth phenotype could result from a decrease in the steady-state levels of the essential Rrp46 protein. To examine this possibility, we assessed the level of each rrp46 protein at both 30°C and 37°C (Fig. 5C) and quantitated results from two independent experiments (Fig. 5D) comparing each rrp46 protein to the level of wild-type protein under the same experimental conditions. This analysis revealed that the only statistically significant change in steady-state protein level detected was for the rrp46-Q86I protein at 30°C (*p* = 0.0307), the fold change in rrp46-L191H at 30°C was not statistically significant (*p* = 0.0710), and the remaining fold changes in rrp46 protein levels at 30°C and 37°C were not statistically significant. This analysis revealed that the level of the yeast rrp46-Q86I variant is decreased compared to wild-type Rrp46 at 30°C where growth defects are not detected. In contrast, the level of the rrp46-L191H protein, which confers growth defects (Fig. 5A,B) when expressed as the only copy of the essential Rrp46 protein, is not reduced at steady-state compared to wild-type Rrp46 at either temperature examined.

To test whether the changes in Rrp46 that model pathogenic mutations present in *EXOSC5* alter the function of the RNA exosome, we examined the processing and levels of well-defined RNA exosome target transcripts in *rrp46* mutant cells (Fig. 6). We analyzed steady-state levels of two precursor transcripts that are processed by the RNA exosome, *U4* snRNA (42) and *TLC1* telomerase RNA (43). The *rrp46-L191H* mutant cells show statistically significant accumulation of the *U4* snRNA precursor (Fig. 6A) as well as the *TLC1* (Fig. 6B) precursor with no significant precursor accumulation observed for the other *rrp46* mutants examined.

**Figure 6.**
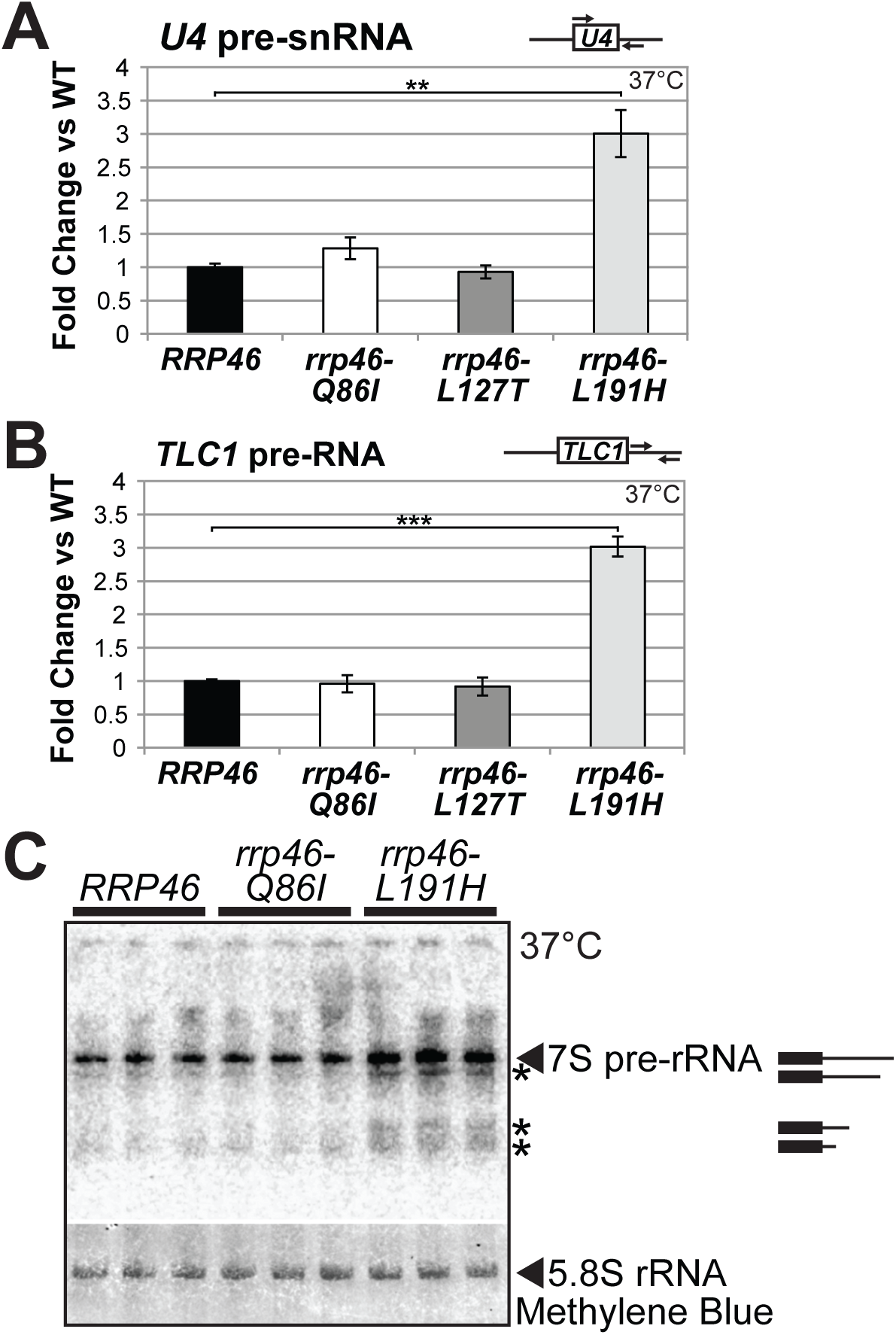
Budding yeast cells expressing the rrp46-L191H variant, corresponding to the EXOSC5 p.Leu206His variant, show impaired processing and elevated levels of some RNA exosome target transcripts at 37°C. (A) The *rrp46-L191H* mutant cells show a statistically significant increase in the levels of non-coding RNA exosome target *U4* snRNA precursor (*p* = 0.0068; very significant (**)) relative to control *RRP46* cells for which the value was set to 1.0 or other *rrp46* mutants analyzed. Total RNA from *rrp46Δ* cells containing only *RRP46, rrp46-Q86I, rrp46L127T*, or *rrp46-L191H* grown at 37°C was reverse transcribed using random hexamers and measured by quantitative PCR using primers indicated in the schematic that detect the U4 snRNA precursor as described in Material and Methods. Relative RNA levels were measured in triplicate biological samples, normalized to a control *ALG9* transcript by the ΔΔCt method, averaged, and graphically shown as fold increase relative to wild-type *RRP46*. Error bars denote standard error of the mean. Statistically significant differences in mean RNA levels were determined using the unpaired t test. (B) The *rrp46-L191H* mutant cells show a statistically significant increase in the levels of non-coding RNA exosome target *TLC1* telomerase precursor RNA (*p* = 0.0008; extremely significant (***)) relative to control *RRP46* cells for which the value was set to 1.0 or other *rrp46* mutants analyzed. Samples were the same as those employed for (A) and were analyzed in the same manner. The location of the primers employed to detect *TLC1* precursor transcript in RT-qPCR is indicated. (C) A northern blot shows that the processing of 7S pre-rRNA to 5.8S rRNA is impaired in *rrp46-L191H* mutant cells, but not in *rrp46-Q86I* mutant cells, compared to wild-type *RRP46* cells. The levels of 7S pre-rRNA and 7S rRNA processing intermediates (asterisks), potentially including 6S pre-rRNA (5.8S rRNA with an additional 5-8 nucleotides at the 3’-end), are elevated in the *rrp46-L191H* cells, but not in *rrp46-Q86I* cells. The northern blot of total RNA from biological triplicates of *RRP46, rrp46-Q86I*, and *rrp46-L191H* cells grown at 37°C was probed with a 5.8S-ITS2 rRNA-specific probe. The northern blot was stained with methylene blue to detect 5.8S rRNA as a loading control.

One of the key evolutionarily conserved functions of the RNA exosome is the 3’-5’ trimming of 7S pre-rRNA to produce mature 5.8S rRNA (44). We performed northern blotting to visualize processing of 7S pre-rRNA for one *rrp46* mutant that does not show obvious growth defects or accumulation of RNA precursors, *rrp46-Q86I*, and the one that shows growth defects and accumulation of RNA precursors, *rrp46-L191H*. As shown in Figure 6C, *rrp46-L191H* mutant cells, but not *rrp46-Q86I* cells, show accumulation of 7S pre-rRNA and 7S rRNA processing intermediates compared to wild-type *RRP46* cells, indicating that pre-rRNA 3’-end processing is impaired in *rrp46-L191H* cells. Combined, these data show that the Rrp46 L191H amino acid substitution, corresponding to the EXOSC5 p.Leu206His substitution identified in Patient 2, impairs the processing function of the RNA exosome in budding yeast.

To extend the analysis of functional consequences of pathogenic amino acid changes to the mammalian EXOSC5 protein, we modeled each pathogenic mutation in mammalian *EXOSC5*. To experimentally test whether Thr114Ile, Met148Thr, or Leu206His alters the steady-state level of the EXOSC5 protein, we expressed Myc-tagged wild-type mouse EXOSC5, EXOSC5 p.Thr114Ile, EXOSC5 p.Met148Thr, and EXOSC5 p.Leu206His in cultured Neuro2A cells and examined the levels of protein by immunoblotting. As shown in Figure 7A, all EXOSC5 proteins are expressed at approximately the same level as wild-type EXOSC5, suggesting that pathogenic amino acid changes in EXOSC5 modeled here do not grossly affect the steady state levels of EXOSC5, at least in this system.

**Figure 7.**
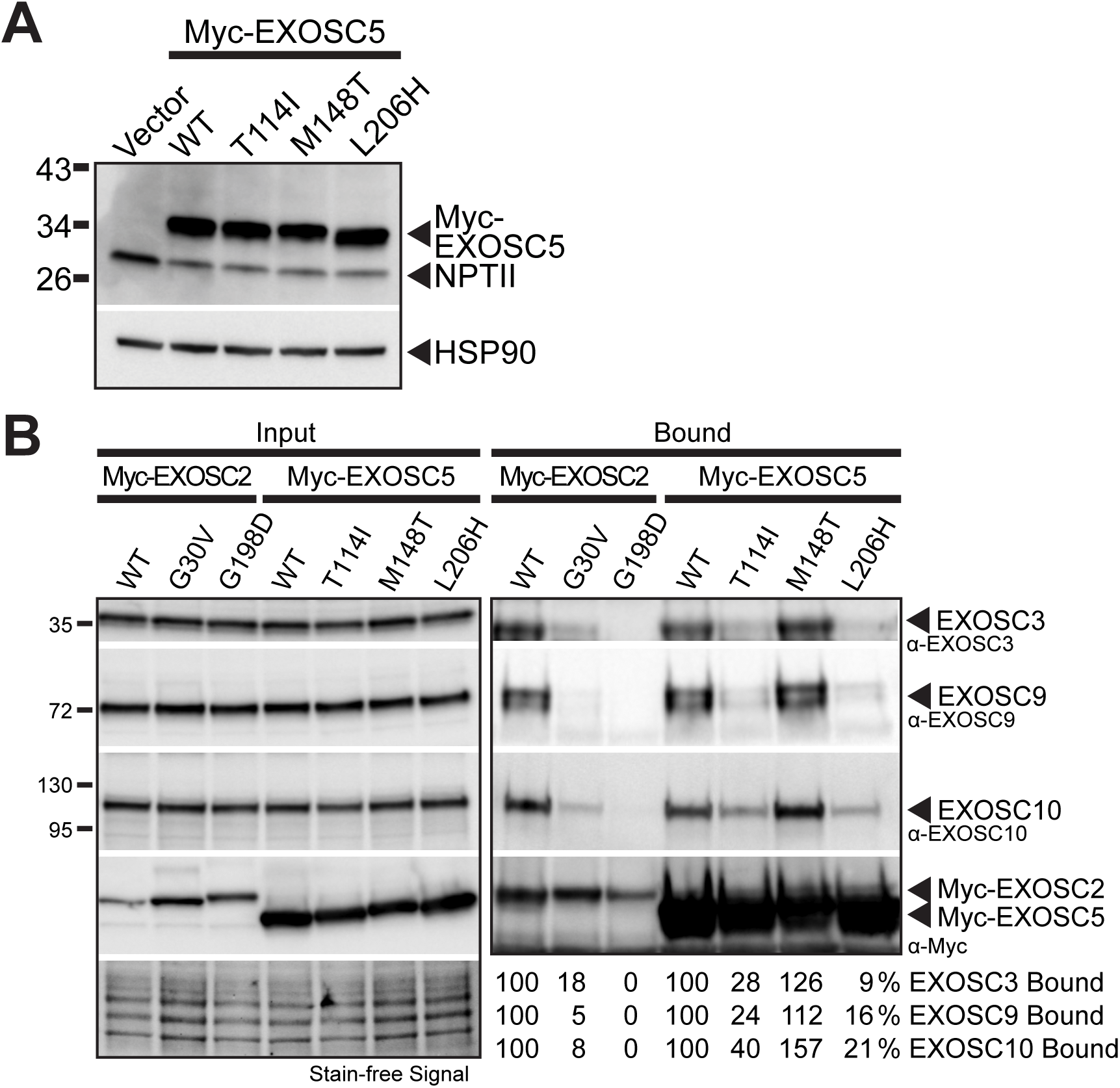
Pathogenic amino acid substitutions in EXOSC5 can alter interactions with other RNA exosome subunits. (A) Murine EXOSC5 variants, corresponding to human EXOSC5 variants identified in patients, are expressed at a similar level to wild-type murine EXOSC5 in a mouse neuronal cell line. The steady-state levels of Myc-tagged murine *EXOSC5* p.T114L (EXOSC5-T114I), *EXOSC5* p.M148T (EXOSC5-M148T), and *EXOSC5* p.L206H (EXOSC5-L206H) variants are similar relative to Myc-tagged wild-type EXOSC5 in mouse N2a cells. Lysates of mouse N2a cells transfected with empty vector or vectors expressing murine Myc-EXOSC5, Myc-EXOSC5-T114I, Myc-EXOSC5-M148T, or Myc-EXOSC5-L206H were analyzed by immunoblotting with anti-Myc antibody to detect Myc-EXOSC5 proteins. HSP90 protein detected with anti-HSP90 antibody serves as a loading control. (A) Murine EXOSC5 variants, corresponding to human EXOSC5 variants identified in patients, show decreased interactions with exosome subunits in a mouse neuronal cell line. Myc-EXOSC5, Myc-EXOSC5-T114I, Myc-EXOSC5-M148T, or Myc-EXOSC5-L206H was immunoprecipitated from N2A cells and interactions with the RNA exosome subunits EXOSC3, EXOSC9, and EXOSC10 were analyzed by immunoblotting. As a control, we also examined the interactions of a pathogenic Myc-EXOSC2-G198D variant, which was recently found to have reduced interactions with RNA exosome subunits (45). Both the Input and Bound samples are shown for Myc-EXOSC2 and Myc-EXOSC5. The fraction of EXOSC3, EXOSC9, or EXOSC10 bound (% Bound) is indicated on the bottom right.

We next employed co-immunoprecipitation to assess interactions between EXOSC5 proteins and other subunits of the RNA exosome complex. Myc-tagged wild-type mouse EXOSC5, EXOSC5 p.Thr114Ile, EXOSC5 p.Met148Thr, or EXOSC5 p.Leu206His was purified from N2A cells and associated EXOSC3 (cap subunit), EXOSC9 (core subunit) or EXOSC10 (catalytic subunit) was analyzed (Fig. 7B). The percentage (%) EXOSC subunit Bound relative to wild-type EXOSC5 is indicated below for each pathogenic EXOSC5 protein. EXOSC5 p. Met148Thr shows interactions with RNA exosome subunits that are comparable to wild-type EXOSC5. In contrast, EXOSC5 p. Leu206His shows only minimal interaction with EXOSC3 (9%), EXOCS9 (16%) or EXOSC10 (21%), while EXOSC5 p.Thr114Ile shows decreased interaction with EXOSC3 (28%), EXOSC9 (24%), and EXOSC10 (40%). As a control, we examined the interactions of a pathogenic EXOSC2 p.G198D, previously shown to have reduced interactions with exosome subunits, and observed similar results (45). These data show that two out of three EXOSC5 protein variants that model pathogenic changes show altered interactions with RNA exosome subunits.

## Discussion

We describe three patients with biallelic variants in *EXOSC5* who share clinical features of delayed development, including impaired motor development and ataxia, and abnormal brain development, with hypoplasia of the cerebellum and brainstem, and ventriculomegaly. All three children showed reduced growth, with heights ranging from the 0-9th centiles. We employed multiple models to establish a requirement for *EXOSC5* in neurodevelopment and define the functional consequences of the amino acid changes present in EXOSC5 in these patients. This analysis reveals that pathogenic mutations in *EXOSC5* can impair the function of the RNA exosome, but different variants likely have distinct mechanistic consequences.

Clinical features common to at least two out of these three, unrelated patients were failure to thrive and short stature, feeding difficulties, developmental delays that predominantly affected motor skills, hypotonia and esotropia (Tables 1 and 2). The brain MRI findings in the first and second patients included hypoplasia of the cerebellum, and in the second and third patients, ventriculomegaly was noted. These findings may comprise a core phenotype associated with biallelic variants in *EXOSC5*. There were also noteworthy differences in the clinical presentations. The first patient had cone-rod dystrophy and complete heart block, whereas the second child exhibited craniosynostosis and pectus carinatum. Of the disease linked to RNA exosome genes, cone-rod dystrophy has been described only with mutations in *EXOSC2* (30), and complete heart block, craniosynostosis and pectus carinatum have not, to our knowledge, previously been linked to RNA exosome subunit variants. Patients 2 and 3 were not evaluated for retinal degeneration, but may have had retinal findings that did not cause readily apparent vision abnormalities, or may have progressed to develop the retinal changes at a later age. We also note that phenotypic variability has been increasingly demonstrated for pathologies linked to the RNA exosome subunit complex genes. For example, two variants, p.Leu14Pro and p.Arg161*, in *EXOSC9* were identified in four patients exhibiting a severe phenotype involving cerebellar hypoplasia, axonal motor neuropathy, hypotonia, feeding difficulties, and respiratory insufficiency consistent with pontocerebellar hypoplasia type intellectual disability (31). The same authors also ascertained two unrelated patients with a milder phenotype resulting from homozygosity for the same missense mutation p.Leu14Pro (31). A similar wide range of phenotypic severity is also probable for variants in *EXOSC5*.

**Table 2.** Clinical features in three patients with biallelic *EXOSC5* variants.

| Clinical feature | Patient 1 | Patient 2 | Patient 3 |
| --- | --- | --- | --- |
| Sex; age at last examination | F; 10 years 10 months | M; 11 months | M; 14 months* |
| Pregnancy | Normal | Normal | Pre-eclampsia |
| Birth | Nuchal cord | Breech position | C-section; 33 w +4 |
| <u>Growth parameters</u> |  |  |  |
| Height | 128.5 cm at 10 y 10 m (2<sup>nd</sup>) | 69.5 cm at 8m 3w (9<sup>th</sup>) | 69.7 cm at 14 m (0<sup>th</sup>) |
| Weight | 31.8 kg at 10 y 10 m (24<sup>th</sup>) | 9.3 kg at 9m 3w (40<sup>th</sup>) | 9.3 kg at 14 m (14<sup>th</sup>) |
| OFC | 52.8 cm at 10 y 10 m (58<sup>th</sup>) | 45.0 cm at 10m 1w (13<sup>th</sup>) | 47.5 cm at 14 m (87<sup>th</sup>) |
| <u>Development</u> |  |  |  |
| Motor | Walked at 8 years | No motor milestones | Mild fine, gross motor delays |
| Speech | Sentences at 10 years | - | Age-appropriate |
| Behavior | Severe OCD | NA | NA |
| <u>Facial anomalies</u> |  |  |  |
| Full cheeks | + | + | - |
| Epicanthus | + | + | - |
| Upslanting palpebral fissures | - | + | - |
| Wide-spaced eyes | - | + | - |
| Broad nasal bridge | - | + | - |
| Micrognathia | + | + | - |
| <u>CNS findings</u> |  |  |  |
| Suck reflex | NA | Absent | NA |
| Grasp reflex | NA | Weak | NA |
| Moro reflex | NA | Weak | NA |
| Limb reflexes | Brisk lower limb reflexes | Reduced | Normal |
| Tone | Mildly reduced truncal tone | Low truncal tone | Normal |
| Gait | Ataxic gait | NA | Normal |
| <u>MRI findings</u> |  |  |  |
| Cerebellar hypoplasia | + | + | - |
| Hypomyelination | - | + | - |
| Small brainstem/pons | + | + | - |
| Enlarged lateral/ 4<sup>th</sup> ventricles | - | + | + |
| Aqueductal stenosis | - | - | + |
| <u>Ocular findings</u> |  |  |  |
| Retinal dystrophy | + | - | - |
| Esotropia | + | + | - |
| Other | Myopia; astigmatism | Persistent pupillary membrane | - |
| <u>Cardiac findings</u> |  |  |  |
| Complete heart block | + | - | Coronary artery fistula |
| <u>Skeletal findings</u> |  |  |  |
| Craniosynostosis | - | + | - |
| Pectus carinatum | - | + | - |
| Hypoplastic T12 ribs | - | Not investigated | + |
| Brachydactyly 5<sup>th</sup> digit | + | - | - |
| Clinodactyly 5<sup>th</sup> digit | + | - | - |
| <u>Muscle biopsy</u> |  |  |  |
| Fiber pathology | Not done | Size variation with hypertrophic and hypotrophic/atrophic fibers | Not done |
| Other | - | Accessory mamillae | Sacral dimple |

M = months; w = weeks; OCD = obsessive compulsive disorder; OFC = occipitofrontal circumference; CNS = central nervous system; MRI = magnetic resonance imaging; NA = not applicable; \*12.5 months corrected chronological age.

**Table 3.**
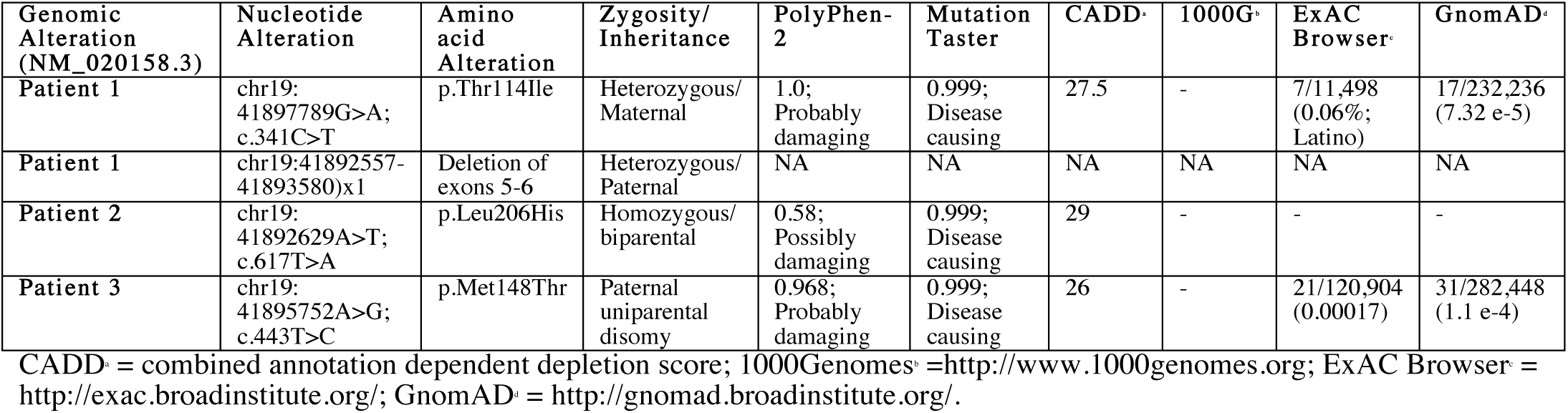
Sequence variants in *EXOSC5*.

Our findings are corroborated by a report of homozygosity for the same p. Thr114Ile variant as seen in heterozygous form in Patient 1, in three individuals who were 27 years, 17 years and 6 years of age and born to second cousin parents of Iranian ethnicity (46). These three family members had mild motor delays and walked at 2-3 years of age with an unbalanced gait. Their intellectual quotients were assessed as ranging between 40 and 50, consistent with moderate intellectual disability (46). All three patients had nystagmus, dysarthria, cerebellar ataxia and increased deep tendon reflexes. A cranial MRI in the 17-year old patient revealed pachygyria and cortical atrophy and ophthalmologic examination of the same patient showed strabismus, but the retina was normal apart from slightly reduced macular reflexes caused by myopia (–2 diopters) (46). Height in the two oldest individuals was measured on the 25^th^ centile and OFCs were all in the normal range (46).

The clinical features reported in the three patients described in the present study overlap with those in prior patients reported to have biallelic, deleterious variants in the genes for exosome subunits already associated with human disease, including *EXOSC3, EXOSC8, EXOSC9* and *EXOSC2* (Table 1). Mutations in these genes cause developmental delays, hypotonia and muscle weakness as typical features (18, 29-32). Mutations in *EXOSC3* and *EXOSC8* cause pontocerebellar hypoplasia, with hypotonia, ataxia, muscle weakness and motor delays and MRI findings of cerebellar hypoplasia, small brainstem and pons and intracerebellar cysts (19, 20, 22, 24, 27, 29, 47). The first patient in our report had ataxia, motor delays and cerebellar hypoplasia, a presentation that has significant overlap with pontocerebellar hypoplasia, but that is milder. Mutations in *EXOSC3* and *EXOSC9* can also cause hypotonia, weakness, motor delays and lower motor neuron signs similar to SMA (18, 32) and the presentation of the second patient in this report overlaps with this constellation of findings. All of the patients with *EXOSC5* variants had a height <10th centile, and failure to thrive and microcephaly have also been observed with mutations in the other exosome subunits (Table 1). With the exception of patients with deleterious *EXOSC2* variants, patients with mutations in RNA exosome subunit genes have typically not had facial dysmorphic features and the *EXOSC5* patients also lacked diagnostic dysmorphic features (Table 1). The third patient had ventriculomegaly and mild motor delays and did not appear to have clinical findings suggestive of pontocerebellar hypoplasia or an SMA-like presentation at an early age. Possibly, his phenotype was influenced by isodisomy for additional autosomal recessive genes. Two female, monochorionic, diamniotic twins have been reported with paternal isodisomy for chromosome 19, who manifest global developmental delays, hypotonia and mild facial anomalies (48). Losses of imprinting for known imprinted genes on chromosome 19 was identified in both these twins, including *ZNF331, PEG3, ZIM2* and *MIMT1*, but there were no deleterious variants on exome sequencing (48). Based on the clinical similarities between patients 1 and 2 and previously reported patients with dysfunction in other RNA exosome subunits and our functional studies, we consider that the variants identified in *EXOSC5* are disease-causing variants. These results add the *EXOSC5* gene of the RNA exosome to the growing number of RNA exosome subunit genes that have been linked to human disease.

The full spectrum of clinical findings that can be associated with deleterious variants in *EXOSC5* and with uniparental isodisomy for chromosome 19 is not yet known and it is possible that features similar to those found in the third patient will be described in other individuals with these molecular abnormalities. Whether a phenotype-genotype correlation will emerge for *EXOSC5* is not yet clear. For *EXOSC3*, homozygosity for p.Asp132Ala results in a milder phenotype with preservation of the pons and the potential for survival into early puberty, whereas compound heterozygosity for *EXOSC3* p.Asp132Ala with an *EXOSC3* nonsense variant or the *EXOSC3* p.Tyr109Asn allele, homozygosity for *EXOSC3* p.Gly31Ala or p.Gly135Glu causes attenuation of the ventral pons and more rapid clinical progression (49). For *EXOSC8*, a phenotype-genotype correlation was suggested for several missense variants, with substitutions such as *EXOSC8* p.Ser272Thr associated with respiratory failure and death in the second year of life, whereas other missense variants were linked to longer survival (49). For *EXOSC9* and *EXOSC2*, only a few families have been reported (30-32), so a comprehensive exploration of phenotype-genotype correlations has not yet been feasible.

As in previous studies of *EXOSC3* (19), *EXOSC8* (29), and *EXOSC9* (32), we employed a loss of function zebrafish model to explore the requirement for *EXOSC5* in neurodevelopment. We found that successful *exosc5* targeting in F_0_ founders resulted in a variety of defects, including mild to moderate tail curvature defects, shortening of the tail and body axis, edema and reduced size and fin defects that resembled the larval phenotypes observed by targeting zebrafish *exosc3, exosc8* and *exosc9* (19, 29, 32). In addition, we observed disruption of midbrain and hindbrain morphology in *mnx1*:EGFP strains that appear to be consistent with the midbrain and cerebellar defects observed with brain imaging of human patients. Our zebrafish model of loss of *exosc5* function thus supports a critical role for *EXOSC5* in brain formation, consistent with past results obtained in zebrafish for loss of function models for other RNA exosome subunits.

To explore how pathogenic variants in the *EXOSC5* gene impact the function of the RNA exosome, we modeled the pathogenic amino acid changes in budding yeast and cultured cells. Taken together, these studies suggest that a decrease in the steady state level of EXOSC5 resulting from pathogenic amino acid substitutions is not likely to explain defects in RNA exosome function. While the steady-state levels of the yeast rrp46-Q86I protein is decreased compared to wild-type Rrp46, neither defects in growth nor RNA processing were detected in these cells. Furthermore, analysis of the steady-state levels of EXOSC5 in cultured mammalian cells revealed no significant changes for the EXOSC5 variants. Recent analysis of EXOSC2 (45) provides evidence that pathogenic variants do not all cause decreased steady state protein levels. Thus, a simple model where pathogenic variants alter protein stability is not sufficient to explain all diseases linked to RNA exosome genes.

We extended the biochemical analysis to explore interactions between EXOSC5 and other RNA exosome subunits. Consistent with the results from modeling *EXOSC5* mutations in budding yeast, this analysis revealed distinct consequences for the pathogenic amino acid changes in EXOSC5. Both the EXOSC5 p.L206H and p.T114I variants show a marked decrease in interaction with other RNA exosome subunits, suggesting impaired assembly or stability of the complex. In contrast, EXOSC5 p.M148T shows no decrease in interactions with RNA exosome subunits. This finding suggests that a subset of the pathogenic variants could alter the overall integrity of the RNA exosome and impair function in this manner. However, these studies do not suggest a mechanistic basis for the altered function of EXOSC5 p.M148T. One possibility is that this amino acid change could alter interaction with critical partners of the RNA exosome such as cofactors. RNA exosome cofactors interact with the complex to confer specificity for RNA targets (50). There is precedence for pathogenic variants in EXOSC3 that were modeled in budding yeast, which to impair interactions with the RNA exosome cofactor, Mpp6 (51). RNA exosome cofactors have been most extensively defined and studied in yeast with work in human cells limited to a few cultured cells lines (50). The possibility exists that RNA exosome cofactors could vary in different cell types to confer specificity for RNA targets that are critical for the specialized functions of those cells. Studies to define the array of RNA exosome cofactors in relevant cell types will need to be undertaken to explore this model and understand whether pathogenic variants could impact these interactions.

Of note, the studies in budding yeast revealed clear functional defects only for rrp46-L191H, which models EXOSC5 p.L206H. While the pathogenic variants lie in conserved regions of EXOSC5, this particular amino acid is the only one strictly conserved between the budding yeast and human proteins (Fig. 4B). The EXOSC5 p.L206H protein also shows the most severe loss of interaction with other subunits of the RNA exosome. Thus, the yeast model system may be valuable to define those pathogenic variants that are most deleterious.

Pathogenic variants in the genes encoding the structural subunits of the RNA exosome, including *EXOSC5*, cause phenotypes that affect specific tissues, such as the brain, despite the ubiquitous expression patterns of these genes (3). EXOSC5 is present on the same face of the RNA exosome core complex as EXOSC3 and directly interacts with EXOSC8 and EXOSC9 (7) (Fig. 4A). The adjacent positions of these subunits and the predicted disturbance to subunit interactions could explain the overlapping clinical presentations between the first two patients and those that have been reported in previously described exosomopathies (Table 1). Mutations in *EXOSC2* have been linked to a unique constellation of clinical findings (30) and the subunit is located on a distinct face in the core complex, suggesting that functional consequences of the missense mutations in this gene may be distinct from those described with the above subunits. The tissue-specific defects could result from variants impacting a limited subset of target RNAs, affecting RNA exosome-mediated degradation/processing of specific noncoding RNAs such as rRNA or tRNA, or by reducing total RNA exosome activity. In addition, the clinical phenotypes show variability amongst the different genes and amongst variants that are predicted to affect related amino acids and subunit interactions across different proteins (3). This variation could be the result of variants that differentially affect the level/stability of the subunit and the integrity of the RNA exosome complex, resulting in variation in the level of functional RNA exosome (3). Indeed, previous studies using patient-derived fibroblasts have shown that pathogenic variants in individual RNA exosome subunit genes can impact the steady-state level of other RNA exosome subunits (32). Alternatively or in addition, pathogenic variants could affect interactions with RNA exosome cofactors that alter the processing/decay of a subset of RNA transcripts. Further mechanistic studies in relevant cell types will be required to define the mechanisms by which pathogenic variants in *EXOSC5* and other RNA exosome genes affect the function of the RNA exosome and cause diverse clinical presentations.

## Conclusion

We describe three patients with delays in motor development, failure to thrive and short stature, feeding difficulties, hypotonia and esotropia and MRI findings including cerebellar hypoplasia and ventriculomegaly. Furthermore, our results demonstrate that different pathogenic *EXOSC5* variants can impair RNA exosome function through distinct mechanisms.

## Materials and Methods

### Case reports

Written, informed consent was obtained from all families for publication of anonymous clinical data. The first and third families provided consent for publication of facial photographs. Summaries of the clinical features for these patients are provided in Table 2. The collaborative study was initiated through the use of the GeneMatcher site (https://www.genematcher.org/) (52, 53).

### Exome sequencing

For patients 1 and 3, clinical exome sequencing (GeneDx, Inc.) was performed as a trio with both biological parents. Using genomic DNA from the proband and parents, the exonic regions and flanking splice junctions of the genome were captured using the Clinical Research Exome kit (Agilent Technologies, Santa Clara, CA). Massively parallel (NextGen) sequencing was done on an Illumina system with 100bp, paired-end reads. Reads were aligned to human genome build GRCh37/UCSC hg19, and analyzed for sequence variants using a custom-developed analysis tool. Additional sequencing technology and variant interpretation protocols have been previously described (54). The general assertion criteria for variant classification are publicly available on the GeneDx ClinVar submission page (http://www.ncbi.nlm.nih.gov/clinvar/submitters/26957). For patient 2, genomic DNA from peripheral blood was used to perform exome sequencing of the patient and both parents. Enrichment was done with SureSelectXT Human All Exon v6 (Agilent Technologies) on 1µg DNA. Alignment was performed with BWA v.0.7.15 (55). PCR-duplicates were marked with Picard (v.1.124) (http://broadinstitute.github.io/picard/); and indel realignment, base quality recalibration, and joint variant calling (HaplotypeCaller) were performed with the Genome Analysis Toolkit [GATK, v. 3.6 (56, 57)]. Functional annotation was performed using Ensembl 85 (58). The variant call files (VCFs) generated were analyzed using the Filtus program (59). Variants with an allelic frequency >0.01 in any of the databases used (gnomAD, ExAC, 1000Genomes, in-house database of >250 whole exomes) were discarded. The analysis was focused on single nucleotide variants (SNVs) and insertion-deletion variants (indels) predicted to be missense, nonsense, frame-shift or splicing variants. Variants with a predicted pathogenicity score <15 according to the Combined Annotation Dependent Depletion (CADD) score (36, 60) were discarded.

### CRISPR/ Cas 9 model of *exosc5* loss of function in zebrafish

All animal experiments were performed under a protocol approved by the Institute for Animal Care and Use Committee (IACUC; AN108657-03) at the University of California, San Francisco. EKW strain zebrafish were maintained at 28°C with a 14 hour (h) light/10 h dark cycle. We designed a CRISPR single guide (sg) RNA targeting exon 2 of *exosc5* (Table S1), the single ortholog of human *EXOSC5*. The sgRNA and Cas9 protein were injected into wild-type eggs from EKW zebrafish at the one cell stage using previously described methods (61-63). We also performed CRISPR/Cas9 injections with eggs from a *mnx1*:EGFP transgenic strain with a 125-bp mnx1 enhancer in a Tol2 vector driving enhanced green fluorescent protein (EGFP) expression in motor neurons (39) and with eggs from an NBT:DsRed strain that labels neurons under the neural ß-tubulin promoter (40). At 24-48 hours post fertilization (hpf), batches of 10-20 larvae were genotyped by Sanger sequencing to assess for successful gene targeting (for primers, see Table S1). All CRISPR/Cas9, F_0_ injected larvae were examined for external abnormalities at 2 to 6 days post fertilization (dpf) using light microscopy, fluorescence microscopy and confocal microscopy (Zeiss LSM 780 NLO FLIM). We examined tail length and curvature, body length and curvature, edema and eye size as previously performed for zebrafish models of *exosc3, exosc8* and *exosc9* loss of function (19, 29, 32, 33). For both typical appearing and deformed larvae, cryosections were made from the heads of larvae at 2 dpf and the corresponding larval tails were genotyped (63). The polymerase chain reaction (PCR) products from larvae with successful *exosc5* targeting were cloned (TOPO® TA Cloning® Kit, Thermo Fisher Scientific) to determine the number of *exosc5* indel variant alleles in each PCR product. Larvae were also phenotyped at 2 dpf using hematoxylin and eosin staining (63). F_0_ larvae with mosaicism for indel variants at the *exosc5* target site were raised in a zebrafish nursery system, but we were not successful in raising F_0_ founders to breeding age.

All procedures involving zebrafish were done in accordance with the NIH guidelines for use and care of live animals and were approved by the UCSF Institutional Animal Care and Use Committee.

### Quantitative RT-PCR in larvae with CRISPR injections targeting *exosc5*

We performed qRT-PCR to determine if there was loss of *exosc5* expression in CRISPR/Cas9 injected larvae compared to wild-type, uninjected controls. RNA was extracted from 2 dpf CRISPR-injected larvae and uninjected control larvae (Thermo Fisher Scientific). cDNA was synthesized using a first strand cDNA synthesis kit (Thermo Fisher Scientific). Ten nanograms of cDNA was amplified using gene-specific primers at final concentrations of 2.5 μM and Universal SybrGreen mastermix containing Rox at a final concentration of 2.5 μM. Reactions were run on a StepOne cycler and analyzed with StepOne™ Software and ExpressionSuite software (Thermo Fisher Scientific) according to the ΔΔCt method. *eef1al1* was used as an internal control gene. Experiments were performed on two independent samples from wild-type larvae and CRISPR-injected larvae from the control EKW strains and each reaction was run in triplicate. Mean RNA levels were calculated by the ΔΔCt method (64) and normalized to mean RNA levels for *eef1al1*. Results were graphed as RNA fold change relative to control with error bars that represent the standard error of the mean (62).

### Protein structure analysis

We used the cryo-EM structure (PDB 6h25) of the human nuclear RNA exosome at 3.8Å resolution (41). Structural modeling was performed using the PyMOL viewer (The PyMOL Molecular Graphics System, Version 2.0 Schrödinger, LLC.). The mCSM webserver was used for predicting the effect of mutations (65).

### *S. cerevisiae* strains and plasmids

The haploid *rrp46Δ* yeast strain (yAV1105) was generated by transformation of a heterozygous diploid *RRP46/rrp46Δ* yeast strain (BY4743 strain background; Open Biosystems, Huntsville, AL) with a wild-type *RRP46 URA3* maintenance plasmid (pAV15; *CEN*) and sporulation of the diploid to yield *rrp46Δ* haploid progeny containing the *RRP46* maintenance plasmid. The wild-type *RRP46 LEU2* plasmid (pAC3482; *CEN*) was generated by PCR amplification of *RRP46* gene containing its endogenous promoter and terminator from yeast genomic DNA using yeast gene specific primers (Integrated DNA Technologies) and cloning into pRS315 (66). The *RRP46-Myc LEU2* plasmid (pAC3485; *CEN*) was generated by PCR amplification of the *RRP46* promoter and coding sequence from yeast genomic DNA using yeast gene specific primers and cloning into pRS315 (66) containing C-terminal 2xMyc tag and *ADH1* terminator. The *rrp46-Q86I LEU2* (pAC3483), *rrp46-L127T LEU2* (pAC3534) and *rrp46-L191H LEU2* (pAC3484) variant plasmids and *rrp46-Q86I-Myc LEU2* (pAC3486), *rrp46-L127T-Myc LEU2* (pAC3535) and *rrp46-L191H-Myc LEU2* (pAC3487) variant plasmids were generated by site-directed mutagenesis of *RRP46 LEU2* (pAC3482) and *RRP46-Myc LEU2* (pAC3485) plasmid, respectively, using oligonucleotides encoding Q86I, L127T, and L191H amino acid changes (Integrated DNA Technologies) and QuickChange II SDM Kit (Agilent). The pcDNA3-Myc-*Exosc5* (pAC3519) plasmid was generated by PCR amplification of the mouse *Exosc5* coding sequence from Neuro-2a cDNA using gene specific primers (Integrated DNA technologies) and cloning into pcDNA3 (Invitrogen) plasmid containing pCMV promoter and N-terminal Myc tag. The pcDNA3-Myc-*Exosc5-T114I* (pAC3520), pcDNA3-*Myc-Exosc5-M148T* (pAC3521), and pcDNA3-*Myc-Exosc5-L206H* (pAC3522) variant plasmids were generated by site-directed mutagenesis of pcDNA3-Myc-*Exosc5* (pAC3519) plasmid using oligonucleotides encoding T114I, M148T, and L206H amino acid changes (Integrated DNA Technologies) and QuickChange II SDM Kit (Agilent). All plasmids were sequenced to ensure the presence of desired mutations and absence of any other mutations.

### *S. cerevisiae* cell growth assays

The *rrp46Δ* yeast strain (yAV1105) was transformed with wild-type *RRP46* (pAC3482), *rrp46-Q86I* (pAC3483), *rrp46-L127T* (pAC3534) or *rrp46-L191H* (pAC3484) variant *LEU2* plasmid and selected on Leu-plates. The Leu+ transformants were grown on 5-flouro-orotic-acid (5-FOA) to select for cells than had lost the *RRP46 URA3* maintenance plasmid (67), and thus only contained the low copy wild-type *RRP46 LEU2* plasmid or *rrp46 LEU2* variant plasmid. Growth of *rrp46Δ* cells containing *RRP46 LEU2, rrp46-Q86I LEU2, rrp46-L127T LEU2* or *rrp46-L191H LEU2* plasmid was assayed on solid media by serially diluting each culture (in 10-fold dilutions), spotting these dilutions on Leu-media and growth at 25°C, 30°C and 37°C. Growth of yeast cells in liquid culture (in rich YPD media) was performed in 96-well format in a shaking and temperature-controlled plate reader (Biotek, Winooski, VT) and optical density (OD600) of the cultures was measured every 10 minutes and plotted. Growth was measured at 30°C and 37°C.

### Immunoblotting

For analysis of C-terminally Myc-tagged wild-type Rrp46 and rrp46 variant protein expression levels, *rrp46Δ* cells expressing Rrp46-Myc (pAC3485), rrp46-Q86I-Myc (pAC3486), rrp46-L127T-Myc (pAC3535) or rrp46-L191H-Myc (pAC3487) protein were grown in 2 ml Leu-minimal media overnight at 30°C to saturation and 10 ml cultures with an OD600 = 0.4 were prepared and grown at 30°C and 37°C for 5 hr. Cell pellets were collected by centrifugation, transferred to 2 ml screw-cap tubes and stored at -80°C. For analysis of EXOSC5 expression levels, mouse N2a cells (68) were transiently transfected with pcDNA3 vector (Invitrogen) containing mouse wild-type *Myc-Exosc5* (pAC3519), *Myc-Exosc5-T114I* variant (pAC3520), *Myc-Exosc5-M148T* variant (pAC3521), *Exosc5-L206H-Myc* (pAC3522) plasmid, or empty vector (pcDNA3; Invitrogen) using Lipofectamine 2000 (Invitrogen) and cells were collected 24 hr after transfection.

Yeast cell lysates were prepared by resuspension of cells in 0.5 ml RIPA-2 Buffer [50 mM Tris-HCl, pH 8; 150 mM NaCl; 0.5% sodium deoxycholate; 1% NP40; 0.1% SDS] supplemented with protease inhibitors [1 mM PMSF; Pierce™ Protease Inhibitors (Thermo Fisher Scientific)], addition of 300 µl glass beads, disruption in a Mini Bead Beater 16 Cell Disrupter (Biospec) for 4 x 1 min at 25°C, and centrifugation at 16,000 x *g* for 20 min at 4°C. Mouse N2a cell lysates were prepared by lysis in RIPA-2 Buffer and centrifugation at 16,000 x *g* for 10 min at 4°C. Protein lysate concentration was determined by Pierce BCA Protein Assay Kit (Life Technologies). Whole cell lysate protein samples (20-50 µg) were resolved on Criterion 4-20% gradient denaturing gels (Bio-Rad), transferred to nitrocellulose membranes (Bio-Rad) and Myc-tagged Rrp46 and EXOSC5 proteins were detected with anti-Myc monoclonal antibody 9B11 (1:2000; Cell Signaling). As loading controls, 3-Phosphoglycerate kinase (Pgk1) protein was detected with anti-Pgk1 monoclonal antibody (1:30,000; Invitrogen) and HSP90 protein was detected with a mouse anti-HSP90 monoclonal antibody (1:2000; Santa Cruz Biotechnology). For transfection control, Neomycin phosphotransferase II (NPTII) expressed from *NeoR* cassette on pcDNA3*-Exosc5* vectors was detected with anti-NPTII monoclonal antibody (1:1000; Cell Applications, Inc.). Primary antibodies were detected using goat secondary antibodies coupled to horseradish peroxidase (1:3000; Jackson ImmunoResearch Inc) and enhanced chemiluminescence signals were captured on a ChemiDoc Imaging System (Bio-Rad).

### Quantitation of immunoblotting

The protein band intensities/areas from immunoblots were quantitated using ImageJ v1.4 software (National Institute of Health, MD; http://rsb.info.nih.gov/ij/) or ImageLab software (Bio-Rad) and mean fold changes in protein levels and percentage protein bound were calculated in Microsoft Excel for Mac 2011 (Microsoft Corporation). To quantitate and graph the mean fold change in rrp46-Myc variant level relative to wild-type Rrp46-Myc level in *rrp46Δ* cells grown at 30°C and 37°C from two immunoblots, R/rrp46-Myc intensity was first normalized to loading control Pgk1 intensity and then normalized to wildtype Rrp46-Myc intensity at 30°C or 37°C for each immunoblot. The mean fold change in R/rrp46-Myc level relative to Rrp46-Myc and standard error of the mean were calculated and graphically represented. The statistical significance of the fold change in each rrp46-Myc variant relative to Rrp46-Myc at 30°C or 37°C was calculated using the Student’s *t*-test on GraphPad software.

To quantitate the percentage of EXOSC3/9/10 bound by wildtype and variant Myc-EXOSC5 and Myc-EXOSC2 from one immunoblot, the EXOSC3/9/10 intensities in the bound lanes on the immunoblot were normalized to the EXOSC3/9/10 intensities in the wildtype Myc-EXOSC5/2 bound lanes and expressed as percentage EXOSC3/9/10 bound. Although the quantitation of EXOSC3/8/9 bound by wildtype and variant Myc-EXOSC5/2 is based on a single immunoblot, the data is representative of multiple experiments.

### Total RNA isolation from *S. cerevisiae*

To prepare *S. cerevisiae* total RNA from cell pellets of 10 ml cultures grown to OD600 = 0.5-0.8, cell pellets in 2 ml screw-cap tubes were resuspended in 1 ml TRIzol (Invitrogen), 300 µl glass beads were added, and cell samples were disrupted in Mini Bead Beater 16 Cell Disrupter (Biospec) for 2 min at 25°C. For each sample, 100 µl of 1-bromo-3-chloropropane (BCP) was added, the sample was vortexed for 15 sec, and incubated at 25°C for 2 min. Sample was centrifuged at 16,300 x *g* for 8 min at 4°C and upper layer was transferred to a fresh microfuge tube. RNA was precipitated with 500 µl isopropanol and sample was vortexed for 10 sec to mix. Total RNA was pelleted by centrifugation at 16,300 x *g* for 8 min at 4°C. RNA pellet was washed with 1 ml of 75% ethanol, centrifuged at 16,300 x *g* for 5 min at 4°C, and air dried for 15 min. Total RNA was resuspended in 50 µl diethylpyrocarbonate (DEPC (Sigma))-treated water and stored at -80°C.

### Quantitative RT-PCR

For analysis of *U4* pre-snRNA and *TLC1* pre-RNA levels in wild-type *RRP46* and *rrp46* mutant cells, *rrp46Δ* (yAV1105) cells containing *RRP46* (pAC3482), *rrp46-Q86I* (pAC3483), *rrp46-L127T* (pAC3534) or *rrp46-L206H* (pAC3484) were grown in biological triplicate in 2 ml Leu-minimal media overnight at 30°C, 10 ml cultures with an OD_600_ = 0.4 were prepared and grown at 37°C for 5 hr. Cells were collected by centrifugation (2,163 x *g*; 16,000 x *g*), transferred to 2 ml screw cap tubes and stored at -80°C. Following total RNA isolation from each cell pellet, 1 µg RNA was reverse transcribed to 1^st^ strand cDNA using the M-MLV Reverse Transcriptase (Invitrogen) and 0.3 µg random hexamers according to manufacturer’s protocol. Quantitative PCR was performed on technical triplicates of cDNA (10 ng) from independent biological triplicates using primers that anneal to the gene body and sequence 3’ of the gene body to detect *U4* precursor (AC5722-5’-AAAGAATGAATATCGGTAATG; AC5723-5’-ATCCTTATGCACGGGAAATACG), and primers that anneal to sequence 3’ of the gene body for *TLC1* (AC7593-5’-GTATTGTAGAAATCGCGCGTAC; AC7594-5’-CCGCCTATCCTCGTCATGAAC)andcontrol*ALG9*primers(AC5067-5’-CACGGATAGTGGCTTTGGTGAACAATTAC;AC5068-5’-TATGATTATCTGGCAGCAGGAAAGAACTTGGG) (0.5 µM) and QuantiTect SYBR Green PCR master mix (Qiagen) on a StepOnePlus Real-Time PCR machine (Applied Biosystems; T_anneal_ -55°C; 44 cycles). The mean RNA levels were calculated by the ΔΔCt method (64), normalized to mean RNA levels in *RRP46* cells, and converted and graphed as RNA fold change relative to *RRP46* with error bars that represent the standard error of the mean.

### Northern blotting

For analysis of 7S pre-rRNA levels and processing intermediates in wild-type *RRP46* and *rrp46* mutant cells, *rrp46Δ* cells containing *RRP46* (pAC3482), *rrp46-Q86I* (pAC3483) or *rrp46-L206H* (pAC3484) were grown in biological triplicate in 2 ml Leu-minimal media overnight at 30°C, 10 ml cultures with an OD_600_ = 0.4 were prepared and grown at 37°C for 5 hr. Cells were collected by centrifugation (2,163 x *g*; 16,000 x *g*), transferred to 2 ml screw cap tubes and stored at -80°C. Total RNA from cells was resolved on an Criterion TBE-Urea polyacrylamide gel (Bio-Rad), blotted to a nylon membrane and membrane was probed with radiolabeled 5.8S-ITS2 rRNA (boundary) oligonucleotide (AC4211/Probe 020-5’-TGAGAAGGAAATGACGCT) to detect 7S pre-rRNA and intermediates and stained with methylene blue stain to visualize 5.8S rRNA as a loading control. Total RNA (5µg) was mixed with equal volume of RNA loading dye (1xTBE; 12% Ficoll; 7M Urea; 0.01 bromophenol blue; 0.02% xylene cyanol) and resolved on 10% TBE-Urea polyacrylamide gel in 1xTBE at 200V for 1.5 hr. RNA was transferred to Hybond™-N+ nylon membrane (Amersham, GE Healthcare) at 15V for 100 min in 1xTBE and cross-linked to membrane with UV light (120,000 µJoules) using UV Stratalinker® 2400 (Stratagene). Membrane was incubated in Rapid-hyb hybridization buffer (Amersham, GE healthcare) at 37°C for 1 hr. DNA oligonucleotide (100 ng) was 5’-end labeled with [γ-P32]-ATP (PerkinElmer) using polynucleotide kinase (New England Biolabs) at 37°C for 30 min. [P32]-Labeled oligonucleotide probe was purified through G25 microspin column (GE Healthcare), heated at 100°C for 5 min, and added to hybridization buffer. Oligonucleotide probe was hybridized to membrane in hybridization buffer at 37°C overnight. Following removal of hybridization buffer, membrane was rinsed twice in 5 x SSPE; 0.1% SDS at 25°C and washed twice in 0.5 x SSPE; 0.1% SDS at 37°C for 20 min each. Membrane was exposed to phosphoscreen overnight and imaged using Typhoon FLA 7000 phosphoimager (GE Healthcare).

### Immunoprecipitation of mammalian RNA exosome subunits

To immunoprecipitate mouse EXOSC5 and EXOSC2, murine Neuro-2a cells were transiently transfected with *Myc-Exosc5* (pAC3519), *Myc-Exosc5-T114I* (pAC3520), *Myc-Exosc5-M148T* (pAC3521), *Myc-Exosc5-L206H* (pAC3522), *Myc-Exosc2* (pAC3463), *Myc-Exosc2-G30V* (pAC3464), or Myc-Exosc2-G198D (pAC3465) plasmid using Lipofectamine 2000 (Invitrogen) and cells were collected 48 hr after transfection. Cells were lysed in HEPES binding buffer [20 mM HEPES, pH 7.2; 150 mM NaCl; 0.2% Triton X-100; Pierce™ Protease Inhibitors (Thermo Fisher Scientific)]. Cell lysates (250-300 µg) in 0.3 ml HEPES binding buffer were incubated with 20 µl Pierce™ anti-c-Myc magnetic beads (Thermo Fisher Scientific) for 3hr at 4°C with mixing. Beads were pelleted by centrifugation, unbound supernatant was removed, and beads were washed three times with 0.3 ml binding buffer. Input (25 µg) and bound Myc bead (1/2 total amount) samples were analyzed by SDS-PAGE and immunoblotting with rabbit anti-Myc monoclonal antibody (1:2000; Cell Signaling) to detect Myc-tagged EXOSC5 and EXOSC2, rabbit anti-EXOSC3 (1:1000; Bethyl Laboratories, Inc.) to detect endogenous EXOSC3, rabbit anti-EXOSC9 (1:1000; Bethyl Laboratories, Inc.) to detect endogenous EXOSC9, rabbit anti-EXOSC10 (1:1000; Bethyl Laboratories, Inc.) to detect endogenous EXOSC10.

## Supplementary Material

*Conflict of interest statement:* YS and GD are employees of GeneDx, Inc.

### Acknowledgements

We thank members of all laboratories who contributed to this work as well as the patients and families. We are grateful to the Diagnostic section at the Department of Medical Genetics at Oslo University Hospital for performing WES on patient 2 and his parents and for providing the VCF files.

