## Supplemental Data for "Biallelic variants in the RNA exosome gene *EXOSC5* are associated with developmental delays, short stature, cerebellar hypoplasia and motor weakness"

Table S1. Primers used in CRISPR/Cas9 experiments

| |
|---|
| Genotyping |
| F: ATTTGATTGGGGTCACGGTA |
| R: CCTTGTACAAACGTGGACGA |
| qRT-PCR |
| F: AGTATCCTGGCTGGAGTTTATG |
| R: CTCGCGGACACTTGGAA |
| Single guide RNA |
| GGGGACACGAGTATCCTGGC |

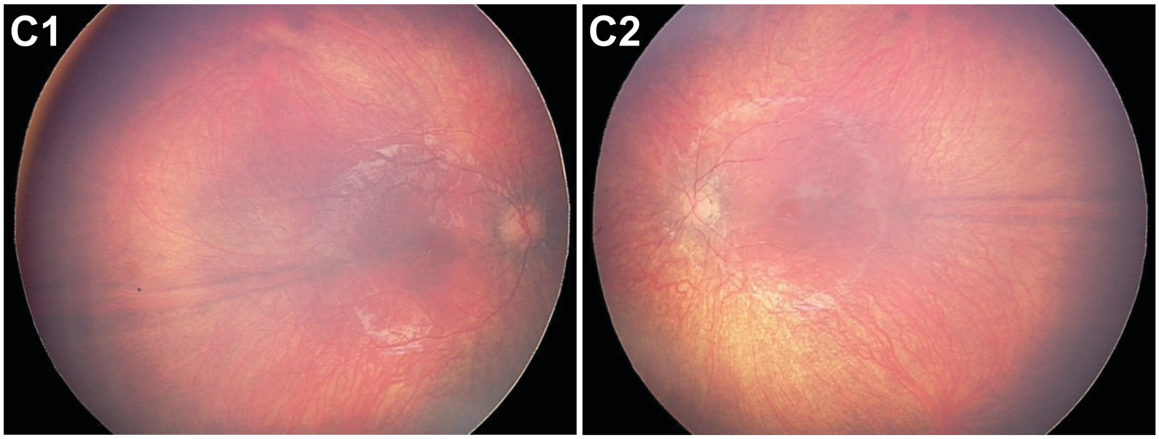


**Figure S1. Color fundus photographs of Patient 1 with biallelic *EXOSC5* variants.**

Color fundus photos of the right (C1) and left (C2) eyes at age 3 years show retinal pigment epithelial mottling along the arcades and mild optic disc pallor in each eye.

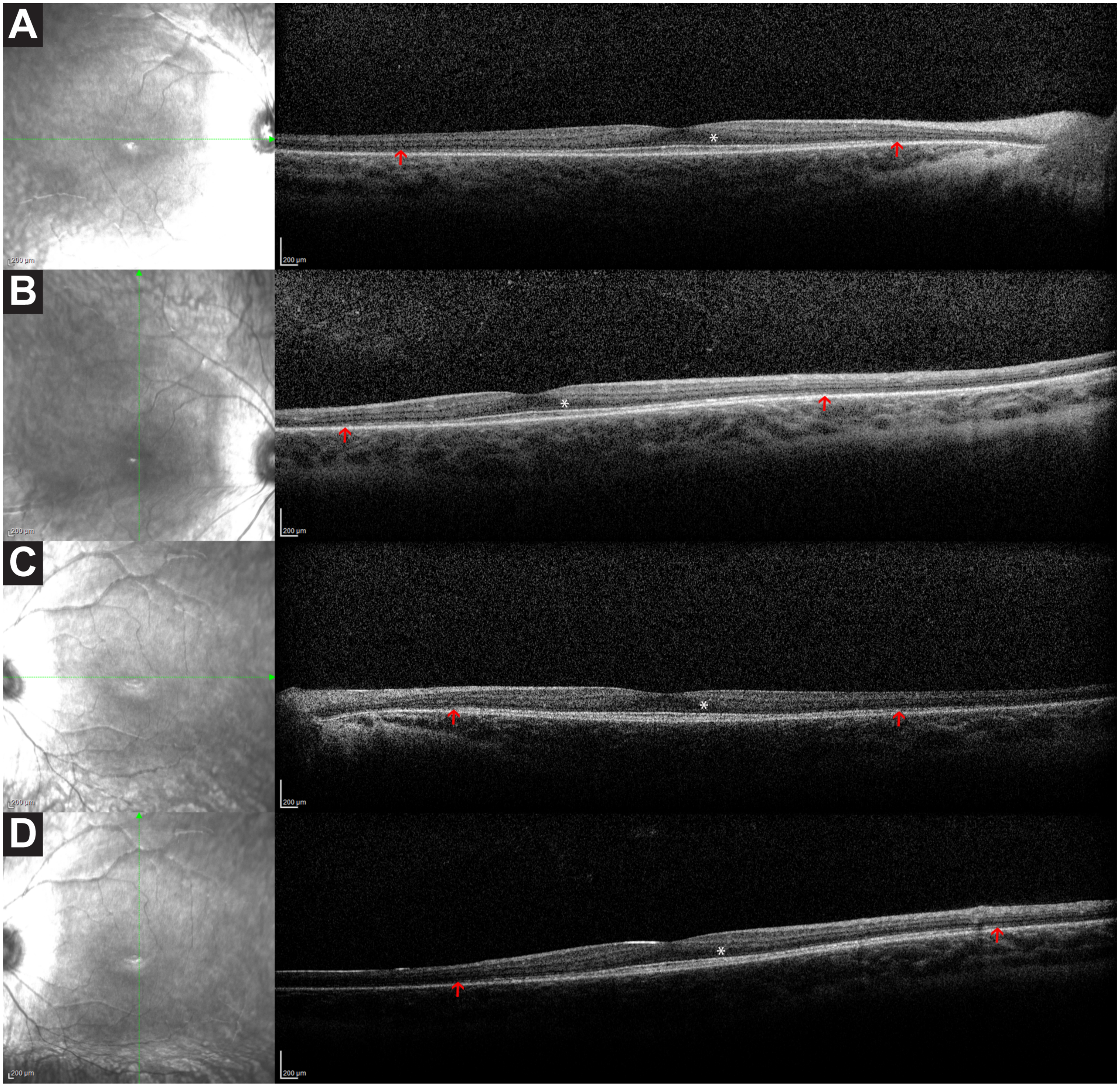


**Figure S2. Spectral domain optical coherence tomograph (SD-OCT) scans of Patient 1 with compound heterozygous biallelic *EXOSC5* variants comprising a deletion of exons 5-6 and a missense variant, p. Thr114Ile.**

Spectral domain optical coherence tomography (SD-OCT) scans of Patient 1 were obtained at age 9 years. The scans extending 30 degrees through the fovea horizontally (A, C) and vertically (B, D) showed thinning of the outer nuclear layer band (*) with loss of the ellipsoid zone band (arrows), representing the photoreceptor inner/outer segment junction, beginning about 10-15 degrees from the foveal center in each scan.

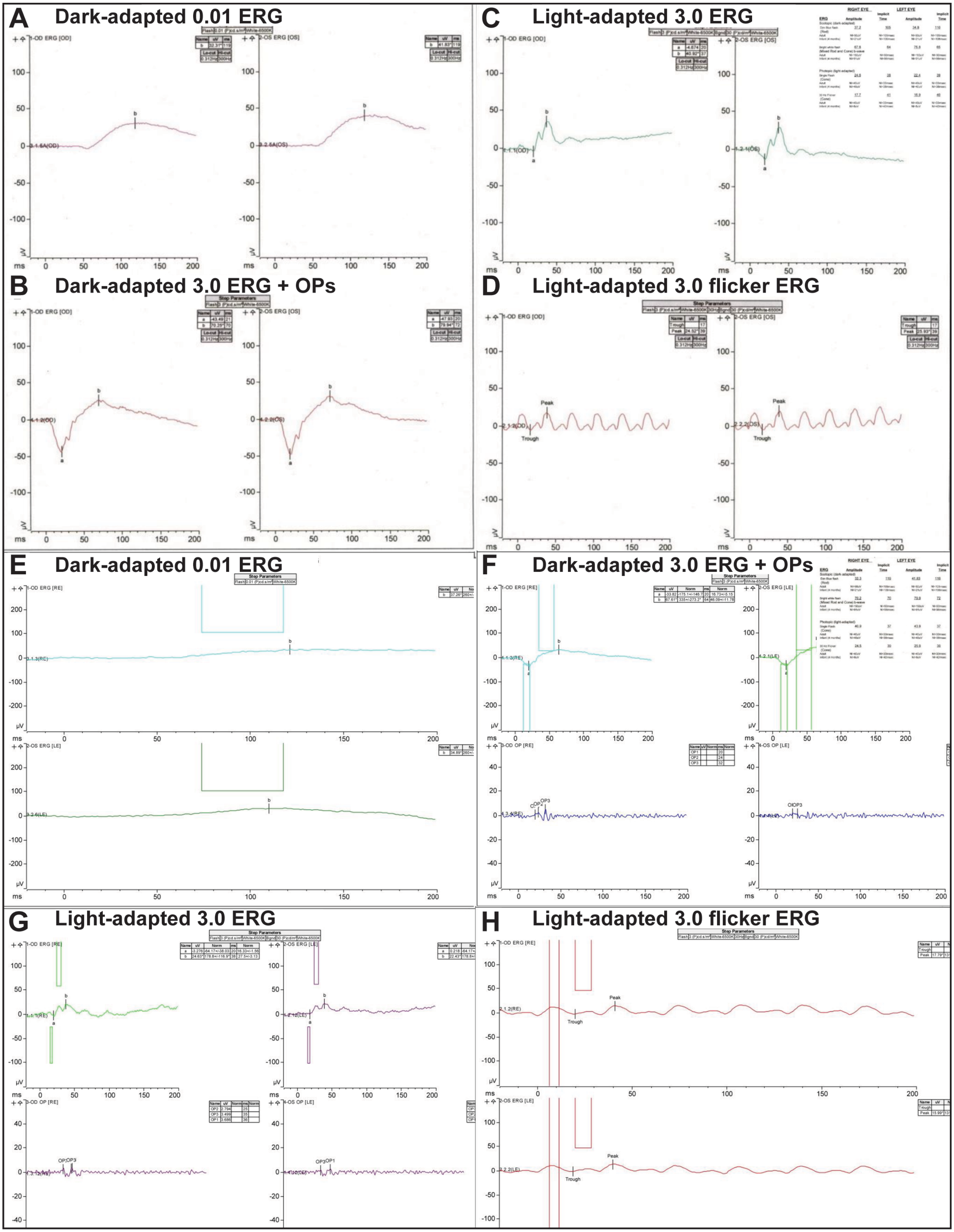
A Dark-adapted 0.01 ERG
B Dark-adapted 3.0 ERG + OPs
C Light-adapted 3.0 ERG
D Light-adapted 3.0 flicker ERG
E Dark-adapted 0.01 ERG
F Dark-adapted 3.0 ERG + OPs
G Light-adapted 3.0 ERG
H Light-adapted 3.0 flicker ERG

Figure S3

**Figure S3. Retinal function in Patient 1 with compound heterozygous biallelic *EXOSC5* variants comprising a deletion of exons 5-6 and a missense variant, p. Thr114Ile.**

Full-field ERG responses recorded according to International Society of Clinical Electrophysiology of Vision standards under brief inhaled anesthesia. (A-D) Scotopic (dark-adapted) (A, B) and photopic (light-adapted) (C, D) responses recorded at age 3 years demonstrated diffuse cone dysfunction to a greater extent than rod dysfunction, manifest as photopic responses reduced below the lower limit of normal by more than scotopic responses. (E-H) Scotopic (E, F) and photopic (G, H) responses recorded at age 6 years showed progressive cone dysfunction, demonstrated as more severely reduced photopic responses compared to scotopic responses. Responses from the right eye are shown on the left and responses from the left eye are shown on the right of panels A-D, F and G. Right eyes are shown at the top and left eyes are shown at the bottom of panels E and H. Insets at upper left of panels (C) and (F) show quantitative measures of amplitudes and timing for each response.

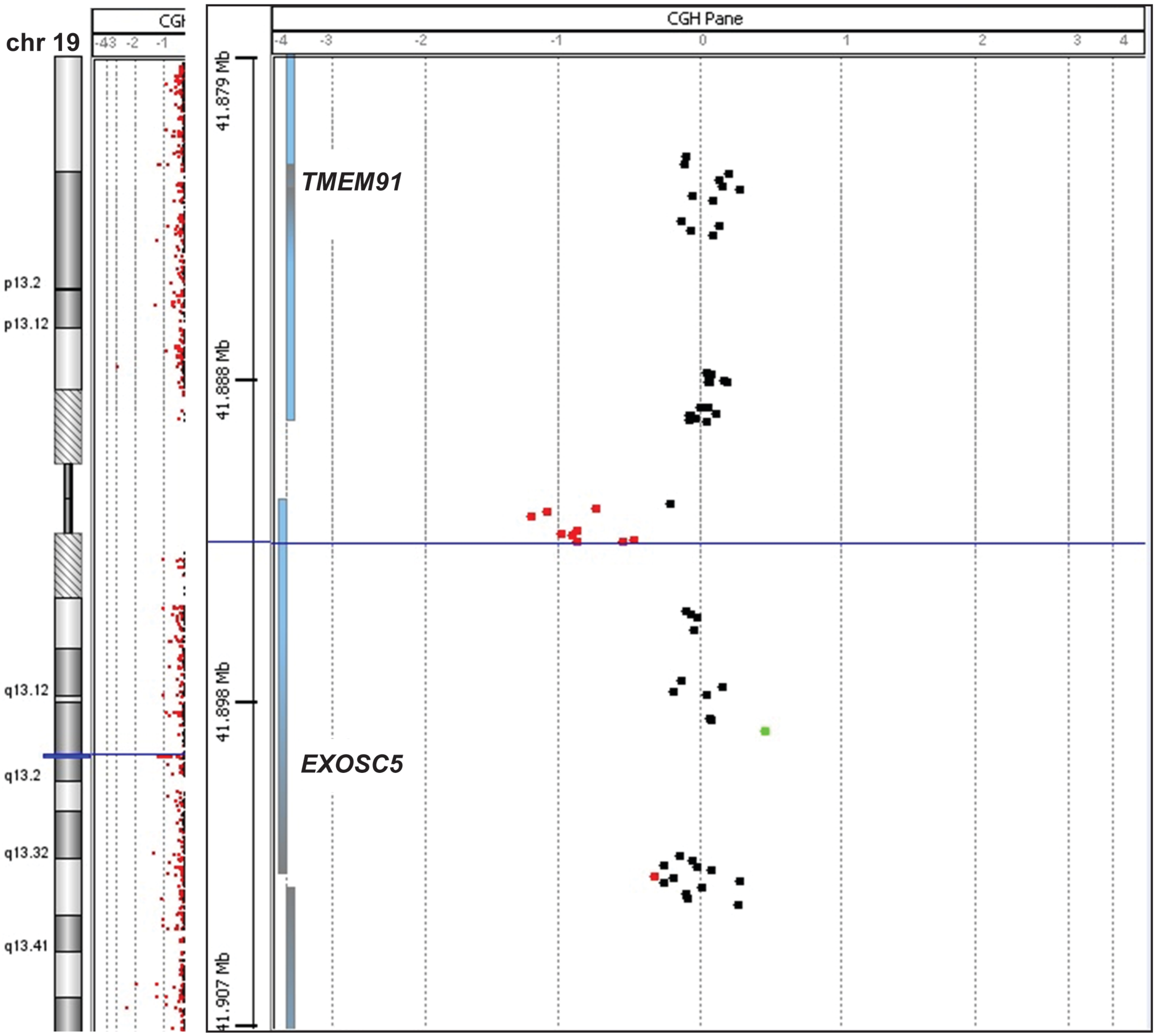


**Figure S4. Exome sequencing in Patient 1 shows a paternally inherited deletion of exons 5-6 in *EXOSC5***

Microarray testing in Patient 1 confirmed a paternally inherited deletion of exons 5-6 in *EXOSC5* initially identified by exome sequencing. The deletion was at least 1,023 bp in size and included exons 5-6 of *EXOSC5*, reported as arr[GRCh37]19q13.2(41,892,557-41,893,580)x1 pat, in Patient 1.

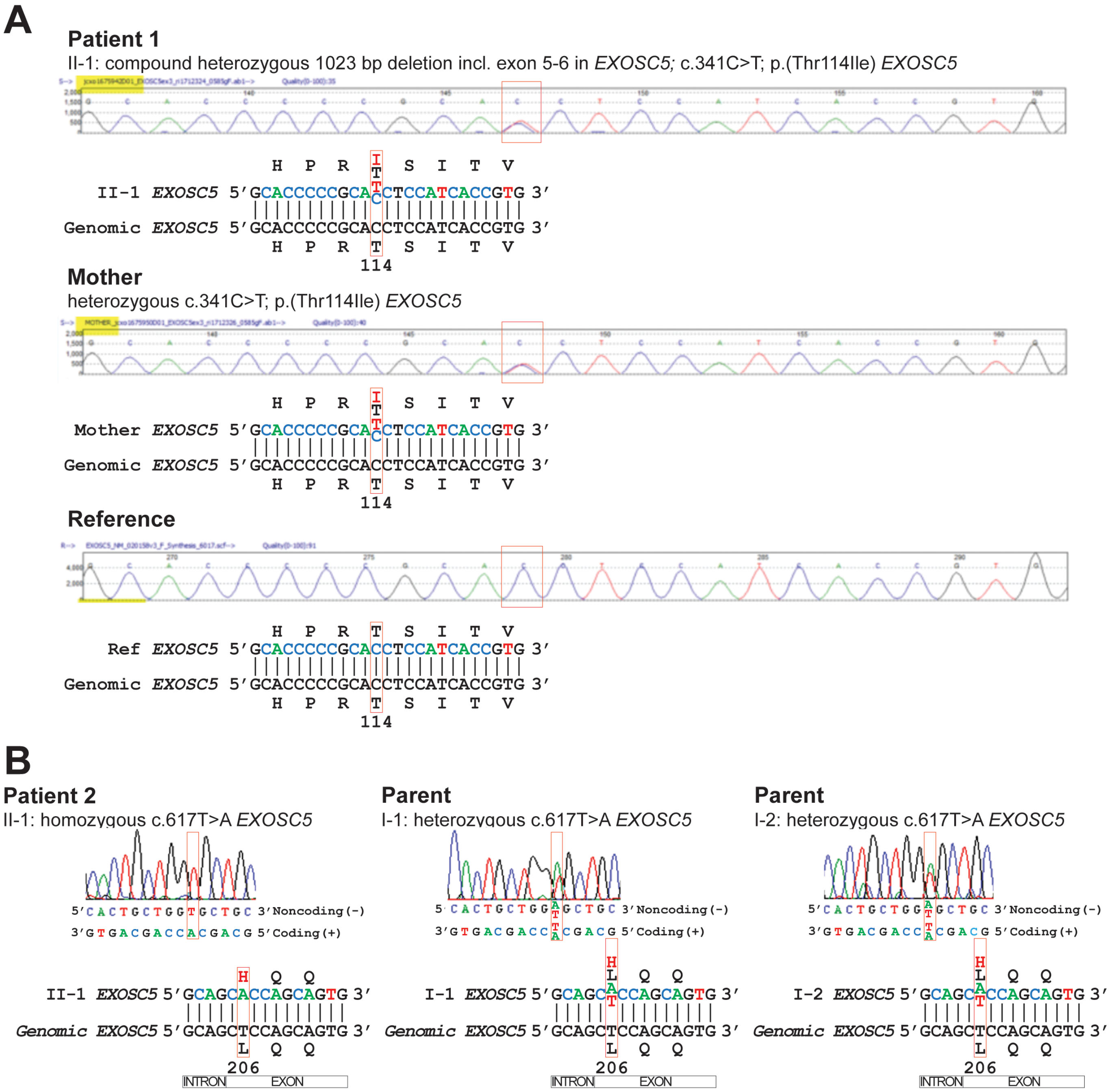


**Figure S5. Sanger sequencing in Patients 1 and 2 showed heterozygous, missense variants in *EXOSC5*.**

(A) Sanger sequencing of the *EXOSC5* gene in Patient 1 confirmed the heterozygous, maternally inherited missense variant, initially identified by exome sequencing: c.341C>T;p.Thr114Ile (chr19(GRCh37):g.41897789;NM_020158.3). (B) Sanger sequencing of the *EXOSC5* gene in Patient 2 confirmed the homozygous missense variant, initially identified by exome sequencing: c.617T>A; p. Leu206His (chr19(GRCh37):g.41892629A>T;NM_020158.3). Sanger sequencing of the *EXOSC5* gene in the parents of Patient 2 confirmed that they are heterozygous for the c.617T>A; p. Leu206His variant.

```
HsapEXOSC5     1 MEEETHTDAKIRAENGTGSSPRGPGCSLRHFACEQNLLSRPDGSASFL-------------------QGDTSVLAGVYGP
Shar           1 METELGG----SAGPDADPGSRSSGCSLRHFACEQNLLSRPDGSASFL-------------------QGDTSVLVGVYGP
Xtro           1 --------------------MEAVGSVLREYGCEQSLLSRPDGSATFL-------------------QGDTSVMAGVYGP
Drer           1 ------MDATSRLCMRTEMELERSCPVLREYGSEQSLLSRPDGSSTFV-------------------QGDTSILAGVYGP
Dmel           1 --------------MNLPAKMDVEKDKLRQMHCEFNPLSRCDGSVMYS-------------------QGATGLIGAVLGP
Cele           1 -----------------------MAGRLREMRCELSFLKNADGSACFS-------------------QGATCIWASCSGP
Cgig           1 --------------------MEPEISKLRQLSSTLGDLSRPDGDATVS-------------------QGDTCVTSAVYGP
Nvec           1 -------------MAAQDNTRPLQTCELRAMFCEHGLLDKADGSSKFA-------------------QGDTQVMAAAYGP
Aque           1 --------------MNIIRHVVTMEVKSRGIECELGIITRADGSAQVK-------------------QGNTIVTCGVYGP
Cowc           1 ------------MVTRAD---GRMPNQLRELSIEHAVLSRADGSSRFS-------------------QRDTSALTAVYGP
Spun           1 ------------MTQRTRPDKRHSPNEMRPFYCSSALLSRADGSVRFK-------------------SGDSAVMCSVFGP
Rdel           1 --------------MVTRPDRRADNKQTRTLSASQNILSRADGSAKFEFGIKRNQTYCYLHAYICLQKGTTSVICSVSGP
Umay           1 ------MVEATSTEGPSESARRTSPTALRSLSAEFGILARSDASASFS-------------------FGPVNVVASVSGP
Cneo           1 MSSY-RTCTTMATAGPSRRPDGRTPAQLRPLHLSIGELDRADGSARFA-------------------FGSNAVLASCSGP
Spom           1 -------------------------------MNRIGILSRSDGSSEWK-------------------QGSARVICGVNGP
Scer           1 ----------------------------MSVQAEIGILDHVDGSSEFV-------------------SQDTKVICSVTGP
Ncra           1 -------------------------MATVTPEASLNALPRADGSARYS-------------------YAGYTVTASVNGP
Afum           1 ---------------------------MVGPTASLTPLKRADGSASYKCP-----------------SSGFDILGSVNAP
Ddis           1 ------------MKTEHTRNDGRCENSIRPVESEQGSLNKADGSAKFS-------------------QDKSKVLAAVYGP
Atha           1 --------------MEIDREDGRTPNQLRPLACSRNILHRPHGSASWS-------------------QGDTKVLAAVYGP
Osat           1 --------------MEESRADGRNPNQLRPFSCTRNPLDRAHGSARWA-------------------QGDTIVLAAVYGP
Ppat           1 MVMAKVEVGAAPSLNVAERADGRSASQLRPLSLSRGLLTRAHGSATWS-------------------QENTTVLAAVYGP
Crei           1 --------------------------MHRTLVCERAVLDRADGSAKWT-------------------QEGSSVLAAVYGP

HsapEXOSC5    62 AEVKVS-KE--IFNKATLEVILRPKIGLPGVAEKSRERLIRNTCEAVVLGT--LHPRTSITVVLQVVSDAG---------
Shar          58 AEVKVS-KE--IFNKATLEVILKPKIGLPGVAEKSRERLIRNTCEAVVLGT--LHPRTSITIALQIVSDAG---------
Xtro          42 AEIKVS-RE--IHNKATLEVILRPKTGLPAIQEKNQEQLIRETCESVIIGS--LHPRTSITIVLQIVSDAG---------
Drer          56 AEVKVS-KE--IYDRATVEVLIQPKMGLPSVRERAREQCVRETCEAALLLT--LHPRSSLTVILQVVHDDG---------
Dmel          48 IEVKTQ-NL--SIDGSYLECNYRPKAGLPQVTDRIREAAIRDVLELALLSE--AHPRSKMSVQIQELEDRG---------
Cele          39 GDVHAS-KA--SDEAMTLDISYRANCGDN--KFNVLNNIIHSTLSNAINLE--LFPHTTISVTVHGIQDDG---------
Cgig          42 GEVKVT-KE--LLDKATIDVVYKPKSGLPGCAEKLLERTIRNSCETVILGN--LHPRSSISITVQEIQNRG---------
Nvec          49 VEVKLN-KE--LIDRATLEVIFRPKIGIPGCSEKLVEGIIRNSCEPIVLTA--LHPRASLTIVVQVVQNSG---------
Aque          48 VEAKSA-RE--KVDKAVVEVIVKSETGLPGPYEKELELLMSSCCETMILTH--LHPHTAISVSIQIQSDNG---------
Cowc          47 AEVKSS-KE--LLDKATVQTVFRPKTGLAGVDDHVCEAILRQALEPVIQRT--ANPRTSITVIVQVMHDDG---------
Spun          50 MEVKLR-DE--KLDKAVVEVIFKTAAGMTTTKERMYERIIRQTLEASILSG--LHPRTSIRVTLQVLTDDG---------
Rdel          67 VEVQMR-DE--KLDEATVEVIVRPAKGVRATKEKLIENTLRTTFEPIILGG--MMPRTLIQITVQVIKDDG---------
Umay          56 TEVRIR-DE--LTDRATLDVIYQPQHGVAGIPARAVSDALTTAFSSVLLLH--HHPRSLIQLVLQTLSSPSLPQSAGQPL
Cneo          61 IEVRLR-EE--LPDKATFEVNHRPLEGVGATPSRALVTTLETIFPPILSLE--KHPRSLVQLVVQSLVPSTGRVVYGSVF
Spom          31 IDVKIR-DE--RLNKATVEVLVQPVSGVAETLEKMISSRIVGILEDAIFLN--TYPRTLIQVSIQIIEEDGT--------
Scer          34 IEPKAR-QE--LPTQLALEIIVRPAKGVATTREKVLEDKLRAVLTPLITRH--CYPRQLCQITCQILESGED--------
Ncra          37 IEAQRR-DE--HAYEAHVDVIVRPSAGVGGTRERHLEFILQSSLSQIILVK--NFPRSVIQIVLQVESTPENAYVNTKLV
Afum          37 VELPGR-RDALKPEEATIEVFVKPGTTPGGVGERYVEGILKTMLGRLILGREKGYPRRGVVLTLAIVGGES-------VC
Ddis          50 IEVNSARKE--KILKSYVEVTFTPAFGNTNYIDKEKELLIKNAVESMILTT--LYPRTQISVILQVFSDDG---------
Atha          48 KAGTKK-NE--NAEKACFEVIWKPKSGQIGKVEKEYEMILKRTIQSICVLT--VNPNTTTSVIIQVVHDDG---------
Osat          48 KPGTRK-GE--NPEKASIEVVWKPMTGQIGKQEKEYEMTLKRTLQSICLLT--VHPNTTTSVILQVVGNDG---------
Ppat          62 KPAAMK-KE--NAERAIIEVVWRAKSGLSGSYEKDAEVVVRRSLEYIILTA--LHPNTAISVILQVINDDG---------
Crei          36 RQAKLQ-KE--DAERAVVEVVFKPRAGLQGHEDRSLELEIRGILEGVIPLG--MFPRTSVMVVLQVLQDDG---------

HsapEXOSC5   128 -----------------------------------------SLLACCLNAACMALVDAGV-PMRALFCGVACALDS----
Shar         124 -----------------------------------------SLLACCLNAACLALVDAGV-PMRALFCGVTCALDP----
Xtro         108 -----------------------------------------SLLSCCLNAACMGLMDAGL-PMRALFCGVTCAMDN----
Drer         122 -----------------------------------------SLLSCCLNAACMALMDAGL-PMSRLFCSVTCAISK----
Dmel         114 -----------------------------------------SIDACAVNCACLAMLIGGL-PLKYSFAAVHAIINE----
Cele         103 -----------------------------------------SMGAVAINGACFALLDNGM-PFETVFCGVLIVRVK----
Cgig         108 -----------------------------------------SLLSCCINSTCMALLDSGV-SMRYLMAAVSCAIND----
Nvec         115 -----------------------------------------SLLSCAVNAACLAMMDAGF-PMRCMMCGITCAITE----
Aque         114 -----------------------------------------SVIVCMVHALCCSLLDACL-PLKTTFSAIKCAFLS----
Cowc         113 -----------------------------------------ALLACALNSASMALIDAGV-PMSAVLAAVTCAWLPAADG
Spun         116 -----------------------------------------GILSTALNAATLALVDAGI-PLRSLAASVTCMIDD----
Rdel         133 -----------------------------------------SVLAASINAIALALLDAGI-PLKYMAAAVTCMFDN----
Umay         131 QTDT------------GDNHRHVP---RQPLLLGPDVPPSATEQAALINAASLALLDAGI-PARASVAACACAILPATQG
Cneo         136 GAEGVGAEQNTWPATDKDDYAYIPESRKDAARISPAAGYTFTARAASINASTLALLSAGTISILALPVAVALVVTT----
Spom          98 -----------------------------------------DTLAAVINGAVLALLDAGI-SLKYIPCAINCHWKNKITQ
Scer         101 -----------------------------------EAEFSLRELSCCINAAFLALVDAGI-ALNSMCASIPIAIIK----
Ncra         112 QA-----------------------------------SLNLPIIPALFQTAVLALLSAAV-PMKATATSTVVAVVSDD--
Afum         109 RG-----------------------------------DSYLTLLPALLHASLLALISASV-PLSMTFAASILAVTS----
Ddis         117 -----------------------------------------SIVSCAINAACLALLDAGI-EMNGLLGSVTLCFNN----
Atha         114 -----------------------------------------SLLPCAINAACAALVDAGI-PMKHLAVAICCCLAE----
Osat         114 -----------------------------------------SLLPCAINACCAALVFAGI-PLKHLAVAIGCGVLE----
Ppat         128 -----------------------------------------SLLACAMNAACAALVDAGI-PLNGLLSAVSCGVTH----
Crei         102 -----------------------------------------GALSCALNAAAAALVDAGV-PLNSMFSSVSCVLTS----
```

```
HsapEXOSC5  162 ------------------------------------------------DGTLVLDPTSKQEKEARAVLTFALDS------
Shar        158 ------------------------------------------------DGALILDPTAKQEKEARAILTFALDS------
Xtro        142 ------------------------------------------------DGTITLDPNFRQQKESRAVLTFAIES------
Drer        156 ------------------------------------------------EGQIITDPTARQEKESRALLTFAIDS------
Dmel        148 ------------------------------------------------QGEYVLDPDQSETLHQRASFTFAFDS------
Cele        137 -------------------------------------------------DELIIDPTAKQEAASTGRVLFSVCKGS----
Cgig        142 ------------------------------------------------NGSIIMDPNTLQEQSTVAMLTFVFEN------
Nvec        149 ------------------------------------------------QDELVLDPTLEQERKATAVLTFVFDS------
Aque        148 ------------------------------------------------DGSMLTDPTTQQEETATSQLTYIIDR------
Cowc        151 S----------------------------------------------SERRLLLDPTLEEQTAATSIATFAFES------
Spun        150 ------------------------------------------------TGELLLDPTAIELERSASVHTFAFDNVSE---
Rdel        167 -----------------------------------------------KTCEVVLDPTAVELQDAKSVHTFAFDNTRK---
Umay        195 LVLRQNSHITEMSQSQLAGLVRQYKQEQDESDDVDRYDEKDSNKRYTAETVVLLDPTPLELDYALSTHVFAFAFSQALDE
Cneo        212 ------------------------------------------------KGRVMLDPEADEEKQAKARLGFGWAWGAVFGT
Spom        136 -------------------------------------------------DEPDVDGTI---NKLESIITICYSISS----
Scer        141 -----------------------------------------------DTSDIIVDPTAEQLKISLSVHTLALEFVN----
Ncra        154 -----------------------------------------------GSKKIVADPSPQDIREATSLHVLAFTS------
Afum        149 ------------------------------------------------SGDIIRQPSVSQAAAAKSLHVLAFSS------
Ddis        151 ------------------------------------------------DGSIYVDPSTEEENQSKAIAIYSFSS------
Atha        148 ------------------------------------------------NGYLVLDPNKLEEKKMTAFAYLVFPNTTLSVL
Osat        148 ------------------------------------------------DGEVILDTNKAEEQQLKSFAHLVFPNSRKSAS
Ppat        162 ------------------------------------------------DGQVFLDPTKPEEQKCKAYVSFVFPSRRLSAV
Crei        136 ------------------------------------------------DRRLVLDPDALEEQAAAARFCFTYPHHFDLTT

HsapEXOSC5  188 ------------------------VERKLLMSSTKGLYSDTELQQCLAAAQAASQ--------------------HVFRF
Shar        184 ------------------------TEQKLLMSTTKGLYSVAEFQQCLAAAQHASH--------------------HIFRF
Xtro        168 ------------------------TERKVLMMSNRGVYSATELQQCIAAAQIASE--------------------KLFQF
Drer        182 ------------------------NERNVLMSSTTGSFSVQELQQCIAISQKASE--------------------QIFQF
Dmel        174 ------------------------VEGNLLLIQTKGSFKIAQFNDIECLCLAASA--------------------EIFQF
Cele        164 -----------------------DGHPEVCAMDAIGHWDFIQLEAAWSLAQPSAS--------------------AIFDF
Cgig        168 ------------------------CNYGVITVSAKGCYTLDQFQQCLSKCRDASK--------------------YVFQF
Nvec        175 ------------------------VNQNLLTSSTKGSFDVDQYNKCLAASKAAMG--------------------NILAF
Aque        174 ------------------------K-GKLVASHATGSFDTEEYIKGLSVASSTAV--------------------HRTDF
Cowc        179 ------------------------TNQEIVFSSTEGQLTVDEYFYTIEMAQKAAA--------------------NVLAF
Spun        179 -------------------------G--TLANMSTGLFTVDEYMRCYETCRLAAV--------------------SVQQL
Rdel        197 -------------------------TSNVLLSDSDGIFNEAEYSSCQEACFEAVG--------------------QIHTF
Umay        275 QD----------------KEATPREISEQIFAESQGAVDLDELQHARRLCLSGCS--------------------AIVAF
Cneo        244 ANEENNM----------GVAGQNDGGAELVWIESEGSFTRQEWSEALQMSKTASK--------------------AILEF
Spom        160 ------------------------EPAKLIFLETAGPIPEEDFFRVLETAPLHAE--------------------EVSKK
Scer        170 ---------------------GGKVVKNVLLLDSNGDFNEDQLFSLLELGEQKCQ--------------------ELVTN
Ncra        181 -------------------------HDELLLSESEGDFTVKEWDGVYENAQKICCQTASKQDGIDMVLDDDASSPDMRHF
Afum        175 -------------------------KGHLLLNESQGTFDFATWEKVHQHASAICRGTLAGNADDDVSMGEEGGEQGLEKF
Ddis        177 ------------------------INKNIVLSKTIGTLTEDQYFKGLDKASESCD--------------------LIISF
Atha        180 P-------------EGSSVAEGEPVEHGIITSITHGVMSVDDYFLCVENGRAATA--------------------SLSAF
Osat        180 S-------------KEP-NQKEEDSERGLITSITHGVMSEEDYFSCIERGLAASS--------------------RISDF
Ppat        194 P-------------ELPADVDGEPVEYGILTSVTRGAMEVEEYFSCVENCRAAAA--------------------KVSEF
Crei        168 AVPAAATGATTASPGADDASATAVVGESVLGSRCVGAFSGDELLDAAALCRRGCE--------------------RVATF

HsapEXOSC5  224 YRESLQRRYSKS--------------------
Shar        220 YRDSLQRRYSKS--------------------
Xtro        204 YRDFIRRRYSKS--------------------
Drer        218 YRDSVKRRYSKMQ-------------------
Dmel        210 YRNQVAKYHGRSEATAKKE-DVNET-------
Cele        201 YKTVMKRKLSVDEQ------------------
Cgig        204 FKDSISKKLSKQ--------------------
Nvec        211 YRQSVERKLSKVSEALILL-SFT---------
Aque        209 YRETISRKMSRIKQ------------------
Cowc        215 MRLSLERKFSKSV-------------------
Spun        212 FRTSSEGKVRREYG------------------
Rdel        232 LRTAVESKKHKEHQQ-----------------
Umay        319 MRTAVQKRVASQVA------------------
Cneo        294 IRIQLDAHLSSHQLS-----------------
Spom        196 MKELLFETYNESDGHENEK-NPKEDVEMDVVA
Scer        209 IRRIIQDNISPRLVV-----------------
Ncra        236 LRSTLETKVASDLHWK----------------
Afum        230 MRETIEDKVYQDYAWKIDAA------------
Ddis        213 IKLAVKNRIMGTNYNENENQNQNEN-------
Atha        227 FRKNFQQSSSKAG-------------------
Osat        226 MRTTLQKQAPGDV-------------------
Ppat        241 SRSSIEQSLKGNRDGC----------------
Crei        228 ARLSLAKSLQAAGGA-----------------
```

**Figure S6. Alignment of EXOSC5 orthologs from diverse animals, fungi and plants.**

Alignment was generated with Clustal Omega. Red and blue letters indicate residues that are identical and similar, respectively, in more than half of the sequences. The residues altered in patients and in yeast are bold and highlighted in yellow. Species included are: *Homo sapiens* (Hsap), *Sarcophilus harrisii* (Shar), *Xenopus tropicalis* (Xtro), *Danio rerio* (Drer), *Drosophila melanogaster* (Dmel), *Caenorhabditis elegans* (Cele), *Crassostrea gigas* (Cgig), *Nematostella vectensis* (Nvec), *Amphimedon queenslandica* (Aque), *Capsaspora owczarzaki* (Cowc), *Spizellomyces punctatus* (Spun), *Rhizopus delemar* (Rdel), *Ustilago maydis* (Umay), *Cryptococcus neoformans* (Cneo), *Schizosaccharomyces pombe* (Spom), *Saccharomyces cerevisiae* (Scer), *Neurospora crassa* (Ncra), *Aspergillus fumigatus* (Afum), *Dictyostelium discoideum* (Ddis), *Arabidopsis thaliana* (Atha), *Oryza sativa* (Osat), *Physcomitrella patens* (Ppat), and *Chlamydomonas reinhardtii* (Crei).

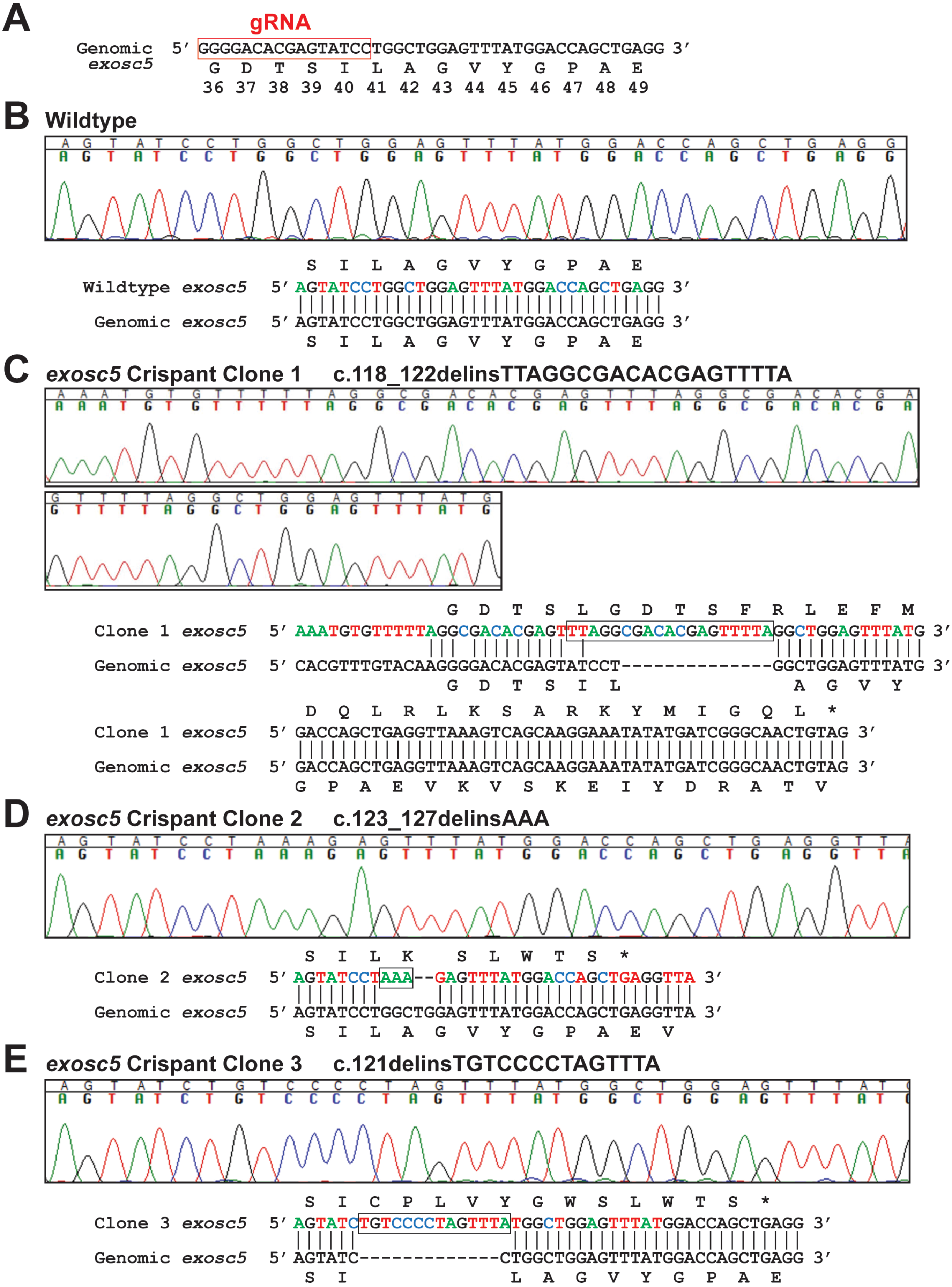
A
gRNA
Genomic exosc5 5′ GGGGACACGAGTATCCTGGCTGGAGTTTATGGACCAGCTGAGG 3′
G D T S I L A G V Y G P A E
36 37 38 39 40 41 42 43 44 45 46 47 48 49
B Wildtype
S I L A G V Y G P A E
Wildtype exosc5 5′ AGTATCCTGGCTGGAGTTTATGGACCAGCTGAGG 3′
Genomic exosc5 5′ AGTATCCTGGCTGGAGTTTATGGACCAGCTGAGG 3′
S I L A G V Y G P A E
C exosc5 Crispant Clone 1 c.118_122delinsTTAGGCGACACGAGTTTTA
G D T S L G D T S F R L E F M
Clone 1 exosc5 5′ AAATGTGTTTTTAGGCGACACGAGTTTAGGCGACACGAGTTTTAGGCTGGAGTTTATG 3′
Genomic exosc5 5′ CACGTTTGTACAAGGGGACACGAGTATCCT--------------GGCTGGAGTTTATG 3′
G D T S I L A G V Y
D Q L R L K S A R K Y M I G Q L *
Clone 1 exosc5 5′ GACCAGCTGAGGTTAAAGTCAGCAAGGAAATATATGATCGGGCAACTGTAG 3′
Genomic exosc5 5′ GACCAGCTGAGGTTAAAGTCAGCAAGGAAATATATGATCGGGCAACTGTAG 3′
G P A E V K V S K E I Y D R A T V
D exosc5 Crispant Clone 2 c.123_127delinsAAA
S I L K S L W T S *
Clone 2 exosc5 5′ AGTATCCTAAA--GAGTTTATGGACCAGCTGAGGTTA 3′
Genomic exosc5 5′ AGTATCCTGGCTGGAGTTTATGGACCAGCTGAGGTTA 3′
S I L A G V Y G P A E V
E exosc5 Crispant Clone 3 c.121delinsTGTCCCCTAGTTTA
S I C P L V Y G W S L W T S *
Clone 3 exosc5 5′ AGTATCTGTCCCCTAGTTTATGGCTGGAGTTTATGGACCAGCTGAGG 3′
Genomic exosc5 5′ AGTATC-------------CTGGCTGGAGTTTATGGACCAGCTGAGG 3′
S I L A G V Y G P A E

Figure S7

**Figure S7. Chromatograms showing cloned *exosc5* indel variants in zebrafish.**

(A) A schematic shows the sequence of the *exosc5* gene in zebrafish with the corresponding protein sequence. The position of the guide RNA (gRNA), which is located in exon 2 of *exosc5*, is indicated in red. (B) The sequence trace from wild-type *exosc5* sequence in zebrafish. (C) Cloned PCR product (Clone 1) showing c.118_122delinsTTAGGCGACACGAGTTTTA, predicting a frame-shift and premature termination codon, indicated by the *. (D) Cloned PCR product (Clone 2) showing c.123_127delinsAAA, predicting a frame-shift and premature termination codon (*). (E) Cloned PCR product (Clone 3) showing c.121delinsTGTCCCCTAGTTTA, predicting a frame-shift and premature termination codon (*).

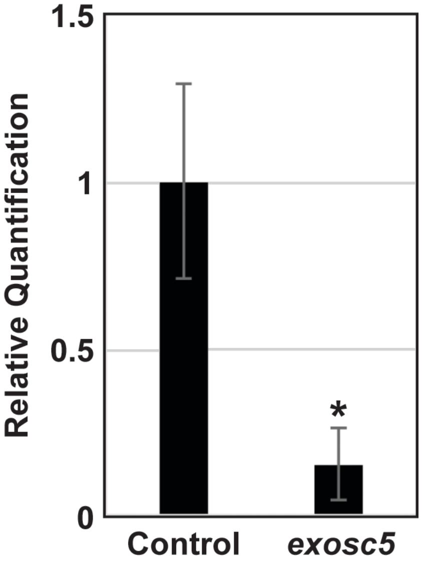


**Figure S8. The *exosc5* expression in *exosc5* crispant zebrafish larvae is significantly reduced compared to control larvae by quantitative RT-PCR.**

RNA was extracted from two biological replicates of EKW zebrafish larvae injected with CRISPR/Cas9 targeting *exosc5* or control EKW zebrafish larvae that were not injected. The cDNA was synthesized with a first strand cDNA synthesis kit (Thermo Fisher Scientific) and then *exosc5* levels were measured by qPCR using an *exosc5* gene-specific primer pair designed to flank the *exosc5* target sequence with one primer crossing an exon-exon junction. After normalization to *eef1al1* as a control gene, *exosc5* expression was significantly reduced (p-value = 0.044 (*)) in CRISPR/Cas9-injected, *exosc5*-targeted larvae compared to uninjected control larvae.
